# Pathogenic MTOR somatic variant causing focal cortical dysplasia drives hyperexcitability *via* overactivation of neuronal GluN2C NMDA receptors

**DOI:** 10.1101/2023.12.01.569539

**Authors:** Louison Pineau, Emmanuelle Buhler, Sarah Tarhini, Sylvian Bauer, Valérie Crepel, Françoise Watrin, Carlos Cardoso, Alfonso Represa, Pierre Szepetowski, Nail Burnashev

## Abstract

**Objective:** Genetic variations in proteins of the mechanistic target of rapamycin (mTOR) pathway cause a spectrum of neurodevelopmental disorders often associated with brain malformations and with intractable epilepsy. The mTORopathies are characterized by hyperactive mTOR pathway and comprise tuberous sclerosis complex (TSC) and focal cortical dysplasia (FCD) type II. How hyperactive mTOR translates into abnormal neuronal activity and hypersynchronous network remains to be better understood. Previously, the role of upregulated GluN2C-containing glutamate-gated NMDA receptors (NMDARs) has been demonstrated for germline defects in the *TSC* genes. Here, we questioned whether this mechanism would expand to other mTORopathies in the different context of a somatic genetic variation of the MTOR protein recurrently found in FCD type II.

**Methods:** We used a rat model of FCD created by *in utero* electroporation of neural progenitors of dorsal telencephalon with expression vectors encoding either the wild-type or the pathogenic MTOR variant (p.S2215F). In this mosaic configuration, patch-clamp whole-cell recordings of the electroporated, spiny stellate neurons and extracellular recordings of the electroporated areas were performed in neocortical slices. Selective inhibitors were used to target mTOR activity and GluN2C- mediated currents.

**Results:** Neurons expressing the mutant protein displayed an excessive activation of GluN2C NMDAR-mediated spontaneous excitatory post-synaptic currents. GluN2C-dependent increase in spontaneous spiking activity was detected in the area of electroporated neurons in the mutant condition and was restricted to a critical time-window between postnatal days P9 and P20.

**Significance:** Somatic MTOR pathogenic variant recurrently found in FCD type II resulted in overactivation of GluN2C-mediated NMDARs in neocortices of rat pups. The related and time-restricted hyperexcitability was sensitive to subunit GluN2C-specific blockade. Our study suggests that GluN2C-related pathomechanisms might be shared in common by mTOR pathway-related cortical dysplasia.

**Key points:**

- Excessive activation of GluN2C NMDAR-mediated currents in spiny stellate neurons expressing FCD-causing MTOR somatic variation
- GluN2C-dependent increase in spontaneous spiking activity in rat somatosensory cortex containing mutant MTOR-expressing neurons
- GluN2C-dependent excessive network activity is time-restricted to a critical period between P9 and P20

## Introduction

The mechanistic target of rapamycin (mTOR) pathway is an intracellular signalling pathway that regulates key cellular processes such as cell growth and metabolism, protein and lipid syntheses, autophagy and many others *via* two protein complexes, namely mTOR complexes 1 (mTORC1) and 2 (mTORC2) (Szwed et al., 2021). In the human brain, excessive, abnormal activation of the mTOR signaling pathway caused by germline or somatic genetic variations results in various epileptic disorders often associated with malformations of cortical development (MCD) and with behavioral (e.g. autistic manifestations) and cognitive impairments (Baulac, 2016; Crino, 2015; Marsan and Baulac, 2018; Nguyen and Bordey, 2021). Collectively designated as mTORopathies, the related disorders include tuberous sclerosis complex (TSC), hemimegalencephaly (HME) and focal cortical dysplasia (FCD) type II. FCD type II are characterized by cortical dyslamination, neuronal heterotopia and the presence of dysmorphic neurons (in FCD types IIa and IIb) and of balloon cells (in FCD type IIb only) (Blumcke et al., 2021b; Mühlebner et al., 2019). All genetic causes of FCD type II identified so far involve members of the mTOR pathway and lead to hyperactivation of mTORC1: hence, single-hit, gain-of-function pathogenic somatic variants were identified in the eponymous MTOR protein that is part of the mTORC1 complex, or in positive mTOR regulators (e.g. the RHEB protein); also, double-hit loss-of-function defects were reported in negative mTOR regulators (e.g. in the TSC proteins hamartin and tuberin, encoded by the *TSC1* and *TSC2* genes, respectively; or in the GATOR1 complex proteins DEPDC5, NPRL2 and NPRL3). Altogether, genetic variants of the mTOR pathway have been identified in 30-60% of FCDs type II (Baldassari et al., 2019; Becker and Beck, 2018).

Upregulation of the mTOR pathway is an important cause of severe epilepsy: indeed, mTORopathies are the most common cause of intractable epilepsy in childhood (Blumcke et al., 2017; Palmini and Holthausen, 2013; Rowland et al., 2012). FCD type II are most often associated with refractory epilepsy. However, pharmacological and surgical managements of the patients remain unsatisfactory. Hence, identification of the pathomechanisms associated with variations of the mTOR pathway is needed. While several cell-autonomous and non cell-autonomous mechanisms have been proposed in various genetic rodent models (Aronica et al. 2023; Nguyen and Bordey, 2021; Represa, 2019), how the numerous genetic variants in different members of the mTOR pathway lead to abnormal neuronal activity and to hypersynchronous network remains to be better understood.

Glutamate-gated NMDA receptors (NMDARs) are instrumental to synchronicity of neuronal networks and NMDARs dysfunction has been associated with a broad spectrum of developmental brain disorders, including epilepsy (Burnashev and Szepetowski, 2015). NMDARs are di- or tri- heterotetramers composed of two obligatory GluN1 subunits and of two other GluN2(A-D)/GluN3(A,B) subunits. Much of functional differences between NMDARs subtypes is attributed to GluN2/3 subunit identity. GluN2/GluN3 subunits show different temporal and spatial expression patterns in the brain; furthermore, NMDARs composition vary depending on cell types and subcellular localization (Hansen et al., 2017). Previously, we showed that neuronal currents mediated by GluN2C-containing NMDARs in the neocortex played a major epileptogenic role in Tsc^+/-^ heterozygous mice as well as in surgically removed tissues not only from TSC patients, but also from genetically-uncharacterized FCD patients (Lozovaya et al., 2014). Slower kinetics and lower sensitivity to Mg^2+^ block of GluN2C-containing NMDARs would sustain increased synaptic integration and hypersynchrony of neuronal networks. While appealing, whether this GluN2C- mediated pathogenic mechanism associated with defects in the hamartin-tuberin protein complex is expandable to more members of the mTOR pathway and to their related disorders, and whether it would also be operating in the mosaic context of the somatic variations seen in FCDs type II, are two important and related questions that have not been addressed yet.

We recently designed a rat model of FCD type II where a pathogenic, recurrent somatic variant (p.S2215F) of the MTOR protein was expressed in a subset of neuronal excitatory cells of the somatosensory cortex by *in utero* electroporation of the appropriate expression vector in neural progenitors of the dorsal telencephalon (Pelorosso et al., 2019). In this model, the affected focal area is composed by electroporated neurons (expressing the mutant MTOR protein) and by non-electroporated (non-mutant) cells. This situation mimics the characteristic cellular mosaicism seen in FCD type II, where MTOR-p.S2215F or other somatic variations in genes of the mTOR pathway arise in a subset of neuronal cells: a subset of the cells in the brain malformation express the mutant isoform of the corresponding protein, while other cells express only the wild-type isoform. In line with previous reports indicating that the recurrent MTOR-p.S2215F variation leads to increased MTOR kinase activity (Grabiner et al., 2014; Mirzaa et al., 2016; Møller et al., 2016; Nakashima et al., 2015), electroporated cells showed increased S6 phosphorylation consistent with mTORC1 overactivation as well as abnormal migration and dysmorphology (Pelorosso et al., 2019). Here, the functional consequences of the pathogenic MTOR-p.S2215F variation on neuronal and network activities and the possible involvement of GluN2C-mediated NMDAR currents in pathogenesis were analysed in the mosaic context of this rat model of mTOR-related FCD-type II.

## Materials and Methods

### Ethical statement

Animal experimentations were performed in accordance with the French legislation and in compliance with the European Communities Council Directives (2010/63/UE). This study was approved under the French department of agriculture and the local veterinary authorities by the Animal Experimentation Ethics Committee (*Comité d’Ethique en Expérimentation Animale)* n°14 under licence APAFIS#23362-2019121712305437 v4.

### In utero electroporation

Wistar rats (Janvier Labs, France) were raised and mated at INMED Post Genomic Platform (PPGI) animal facility and kept at controlled temperature (21-23°C) under conditions of optimal health hygiene with food and water ad libitum, and 12-12 h dark-light cycle (light period from 8 am to 8 pm). *In utero* injections and electroporations were performed from timed pregnant rats at embryonic day 15.5 (E15.5) as previously described (Pelorosso et al., 2019; Salmi et al., 2013).

Plasmid electroporations were performed with either wild type (WT) or mutant MTOR expression vectors (pCAG-MTOR WT-IRES-GFP or pCAG-MTOR p.S2215F, respectively, 3.0 μg/μL each), or with non-recombinant vector (pCAG-IRES-GFP; 0.5 μg/μL).

### Single cell patch-clamp recordings

Brains were rapidly removed from pups and placed in oxygenated ice-cold protecting solution containing (in mM): 126 choline chloride, 2.5KCl, 1.25 NaH2PO4, 7 MgCl2, 0.5 CaCl2, 26 NaHCO3, 10 D-glucose. 300 μm-thick coronal neocortical slices (Vibratom Leica VT1000S, Leica Microsystems Inc., USA) were then kept for at least 1h in artificial cerebrospinal fluid solution ACSF-1 containing (in mM): 125 NaCl, 3.5 KCl, 1 CaCl2, 2 MgCl2, 1.25 NaH2PO4, 26 NaHCO3 and 10 glucose, equilibrated at pH 7.3 with 95% O2 and 5% CO2 at room temperature (22-25°C).

Whole-cell patch-clamp recordings were performed at room temperature (22-25°C) by using an EPC-10 amplifier and Patch Master software (HEKA Elektronik, Germany) and filtered using a low pass 2 kHz filter. For recordings, slices were pre-incubated in oxygenated Mg^2+^-free ACSF (ACSF-2: same as ACSF-1 but with 2.5 mM CaCl2 and no MgCl2) and then transferred to the recording chamber and perfused with oxygenated ACSF-2 recording solution at 3 mL.min^-1^. Neurons were visualized using an Olympus BX61WI microscope (Olympus, Life Sciences Solutions, USA). Recordings were made in somatosensory cortex of electroporated (GFP^+^) and their adjacent (at < 20 Om distance) non-electroporated (GFP^-^) cells that displayed typical electrophysiological and morphological properties of spiny stellate neurons, which represent the majority (58-77%) of all excitatory neurons in layer 4 of somatosensory cortex in rodents (Scala et al., 2019; Staiger et al., 2004). Patch pipettes were pulled from borosilicate glass capillaries (World Precision Instruments, USA) and had resistance of 4-7 MΩ when filled with the internal solution of the following composition (in mM): 130 K-gluconate, 10 Na-gluconate, 4 NaCl, 4 MgATP, 4 phosphocreatine, 10 HEPES and 0.3 GTP (pH 7.3 with KOH). Biocytin (final concentration 0.3-0.5 %) was added to the solution to label the neurons from which recordings were obtained.

### Action potential (AP) firing pattern and intrinsic properties

Current clamp configuration was used to record the AP firing pattern and intrinsic properties of GFP positive and GFP negative adjacent neurons. For quantification of intrinsic properties, current step recordings (20 steps of 10 pA each from -100 pA to 100 pA) were performed using Patchmaster software (HEKA Elektronik, Germany). Analysis was performed using Clampfit software 11.2.1 (Molecular Devices, USA). The input resistance and membrane capacitance were estimated as done previously (Golowasch et al., 2009).

### Spontaneous excitatory post-synaptic currents recordings and analysis

Spontaneous excitatory post-synaptic currents (sEPSCs) were recorded in voltage clamp mode using Axoscope (Molecular Devices, USA) and Digidata 1322A (Molecular Devices, USA).

All recordings were done in ACSF-2 without or with 10 μM DQP 1105 (Tocris Bio-Techne, France), 10 μM DQP 1105 + 1μM Ro 25-6981 (Tocris Bio-Techne, France), or 50 μM APV (Hello Bio, UK) at room temerature (22-25°C). Membrane holding potential was set at -75 mV (reversal potential of GABAergic currents) to exclude contribution of GABAR-mediated currents. sEPSCs were manually selected from the recordings using MiniAnalysis 6.0.3 software (Synaptosoft Inc., USA). The threshold amplitude for detecting sEPSCs was set at twice the baseline noise (root mean square).

Events that did not show a typical synaptic waveform were rejected. Only events that did not show any sign of multiple peaks (that is, contamination of rise or decay phases by subsequent events) were selected for analysis of amplitude and charge transfer. Averaged traces of sEPSCs were obtained from MiniAnalysis software and further analysed with Origin software (MicroCal, USA). For each neuron, original traces from individual experiments were aligned based on the starts of their rising phases and averaged. These averaged traces from individual experiments were then averaged to form grand average traces. To quantify the current decay kinetics, we measured charge transfer of sEPSCs normalized to the peak amplitude. Normalized charge transfer was calculated by integrating the area under the curve of the current trace between the peak of sEPSC and 200 ms after the peak for each individual cell. Rapamycin-based experiments were performed as described previously (Lozovaya et al., 2014) with daily intraperitoneal injections (3 mg/kg) during seven consecutive days (P8-P15) prior to the recordings; in these experiments, the rapamycin-untreated condition consisted in sequential injections of only the vehicle (NaCl 0.9%, Ethanol 4%, Tween-80^®^ 5%, polyethylene glycol 5%). In some experiments spontaneous bursts of action potential (AP) were recorded in current clamp mode in whole-cell configuration using a Digidata 1322A and Axoscope (Molecular Devices, San Jose, CA, USA). Event detection was performed using MiniAnalysis 6.0.3 software (SynaptosoftInc., Decatur, GA, USA). Briefly, bursts were detected based on the action potentials waveforms. Traces of individual events were then extracted and analysed further on Origin software (MicroCal, Northampton, MA, USA). Duration and area under curve were compared between the start of the event and the time the trace reached the baseline after the last spike.

### Extracellular recordings

For extracellular recordings, slices from P9, P14-P16 and P20-P21 pups of both sexes were placed in a perfused chamber at room temperature (22-25°C) in oxygenated Mg^2+^ containing ACSF (ACSF-1) medium under a Leica DMLFS microscope (Leica Microsystems Inc., USA). The recordings were performed using a coated nichrome electrode with an Iso-DAM8A World Precision Instruments amplifier (World Precision Instruments, USA). Data was digitized using a Digidata 1322A and recorded using Axoscope (Molecular Devices, USA). Spontaneous activity was recorded without or with 10 μM DQP 1105 (Tocris Bio-Techne, France) or 50 μM APV (Hello Bio, UK).

Files were analysed using Clampfit (Molecular Devices, USA). Recordings were filtered with a low pass 1kHz filter and AP frequency was estimated using the threshold event detection with threshold set as twice the background noise. Rapamycin-based experiments were performed as described above.

### EEG recordings

Cortical electroencephalography (EEG) was performed on postnatal days P60-P70 in freely moving MTOR-p.S2215F and MTOR-WT electroporated rats and analyzed as done previously (Sahu et al., 2019) and as further described in Figure S1.

### Histological reconstruction of neurons

A subset of whole-cell recorded neurons were filled with biocytin. After 24 h in paraformaldehyde 3% at 4°C, the sections were then rinsed in PBS and pre-incubated for 1 h in 0.3% Triton X-100 in PBS with 5% NGS (newborn goat serum) at room temperature. Slices were incubated in Streptavidin-Alexa 555 (Invitrogen) in PBS Triton X-100 0.3% and NGS 5% overnight at 4°C. After thorough rinsing, slices were incubated for 10 min with Hoescht 1/1000 in PBS Triton X-100 (0.3%). Finally, slices were mounted in Fluoromount (Thermo Fisher Scientific) and coverslipped. Slices were imaged using confocal microscopy (SP5-X Leica, Leica Microsystems Inc., USA) and neurons were reconstructed using Neurolucida software (MBF Bioscience, USA).

### Immunohistochemistry

Pups (P14-P16) were anesthetized using Zoletil/Domitor (0.4 mg/kg) and intracardiacally perfused with paraformaldehyde (PFA) 4%. Brains were dissected and post-fixed in PFA 4% for 48h. After rinsing with PBS, brain coronal slices (80 μm) were cut using a vibratome (Leica VT1000S; Leica Microsystems Inc., USA). Slices were placed in PBS with 0.1% sodium-azide until immunostaining. Ror-beta and GFP immunostaining was performed as follows: slices were incubated in 10mM Citrate 0.05%Tween, pH=6 for 30 min at 95°C for Ror-beta antigen retrieval.

After cooling down, sections were rinsed several times with PBS. Sections were blocked using PBS with 0.3% Triton X-100, 2% BSA and 3% Goat Serum for 1 h at room temperature under small agitation. Sections were then incubated in PBS with 0.3% Triton X-100, 1% BSA and 1% goat serum overnight at 4°C with mouse anti-Ror-beta primary antibody (1/250, Aves, CA, US) and chicken anti-GFP primary antibody (1/500, Aves, CA, US) for experiments at P14-P16. Sections were washed in PBS and incubated for 2 h at room temperature with a goat anti-chicken Alexa 488 secondary antibody (#A11039, 1/500, Invitrogen, MA, US) and goat anti-mouse Alexa 555 (#A21422, 1/500, Invitrogen, MA, US). Slices were counterstained with Hoescht 33342 (1/1000, Thermo Fisher Scientific, MA, US), washed in PBS and mounted using Fluoromount (Thermo Fisher Scientific, MA, US). Images were acquired using LSm-800 Zeiss confocal microscope (Zeiss, Germany) and treated using ImageJ software.

### Statistics

Comparison of groups was done with Mann-Whitney test (data sets = 2) or with Kruskal– Wallis test followed by Dunn’s correction for multiple testing (data sets > 2). Friedman test with Dunn’s post-hoc correction was used to analyse the effect of DQP 1105 and APV on the frequency of action potentials in MTOR-p.S2215F extracellular recordings. Effects of drugs on sEPSCs area was analyzed with a mixed-effect model followed by Tukey’s multiple comparison test. For all comparisons, sex ratio did not differ significantly between conditions (Fisher’s exact test). All values are given as mean ± SD unless stated otherwise. Level of significance was set at p<0.05. Raw data are provided as Supplementary material (Table S1).

## Results

### Mutant MTOR-p.S2215F neurons display decreased cell input resistance and increased capacitance

We used a model of FCD created by *in utero* electroporation of vectors allowing for the co-expression of GFP with either of the mutant MTOR-p.S2215F or the wild type (WT) MTOR isoforms in neural progenitors from the somatosensory cortex of rat embryos at E15.5. In this model, the presence of neuronal dyslamination (Pelorosso et al., 2019) and of epileptic activity, as assessed by telemetric EEG in rats aged P60 (Figure S1) and as also very recently reported in the corresponding mouse model (Okoh et al., 2023), was confirmed. To start in studying the functional consequences of the pathogenic MTOR-p.S2215F variation, we first studied the activity of neurons expressing MTOR-p.S2215F compared to neurons expressing MTOR-WT using patch clamp recordings in brain slices from P14-16 rats previously electroporated *in utero*. This developmental period was investigated as the immature brain is more susceptible to epileptic activity, and, more precisely, as it corresponded to the critical time window when GluN2C-containing NMDARs played a major epileptogenic role in Tsc^+/-^ heterozygous mice (Lozovaya et al., 2014). We also electroporated the pCAGIG-GFP construct that only encodes GFP (GFP-only) as a negative control. Only recordings from spiny stellate neurons, in which overexpressed GluN2C-mediated currents were detected in TSC (Lozovaya et al., 2014), were included for analyses, based on their distinct firing properties as compared to pyramidal cells (see also Material & Methods section). Analysis of the firing pattern elicited by depolarizing steps revealed that MTOR-p.S2215F neurons displayed significantly decreased input resistance (Rinput) and increased capacitance (Cm) compared to MTOR- WT neurons and to GFP-only neurons (MTOR-p.S2215F: Rinput = 241.3 ± 69.98 MΩ, Cm = 246.2 ± 106.2 pF; MTOR-WT: Rinput = 529.5 ± 276.3 MΩ, Cm = 80.1 ± 46.67 pF, GFP: Rinput = 413.9 ± 120 MΩ, Cm = 75.16 ± 20.83 pF) while no significant difference was detected between the MTOR-WT and the GFP-only neurons ( MTOR-p.S2215F vs MTOR-WT: pRinput = 0.0006 and pCm < 0.0001; MTOR-p.S2215F vs GFP: pRinput = 0.0169 and pCm < 0.0001; MTOR-WT vs GFP: pRinput >0.9999 and pCm = 0.8842; Kurskall-Wallis test with Dunn’s post-hoc test) (Figure 1). Consistent with the altered input resistance and capacitance as detected in MTOR-p.S2215F neurons, and with the previously reported increased cell size of MTOR-p.S2215F neurons (Pelorosso et al., 2019), post-hoc reconstruction of biocytin-filled recorded cells at P14-P16 revealed significant increase of the soma volume and dendrite surface area of MTOR-p.S2215F neurons, as compared to MTOR-WT neurons (Figure S2). No significant change in other neuronal intrinsic properties (i.e. resting potential, rheobase, first action potential (AP) amplitude, first AP halfwidth and AP threshold) was detected (Figure S3). Of note, non electroporated, GFP^-^ neurons adjacent to electroporated, GFP^+^ mutant MTOR-p.S2215F neurons displayed unaltered input resistance and membrane capacitance, as for neurons electroporated with MTOR-WT (Figure S4).

**Figure 1.**
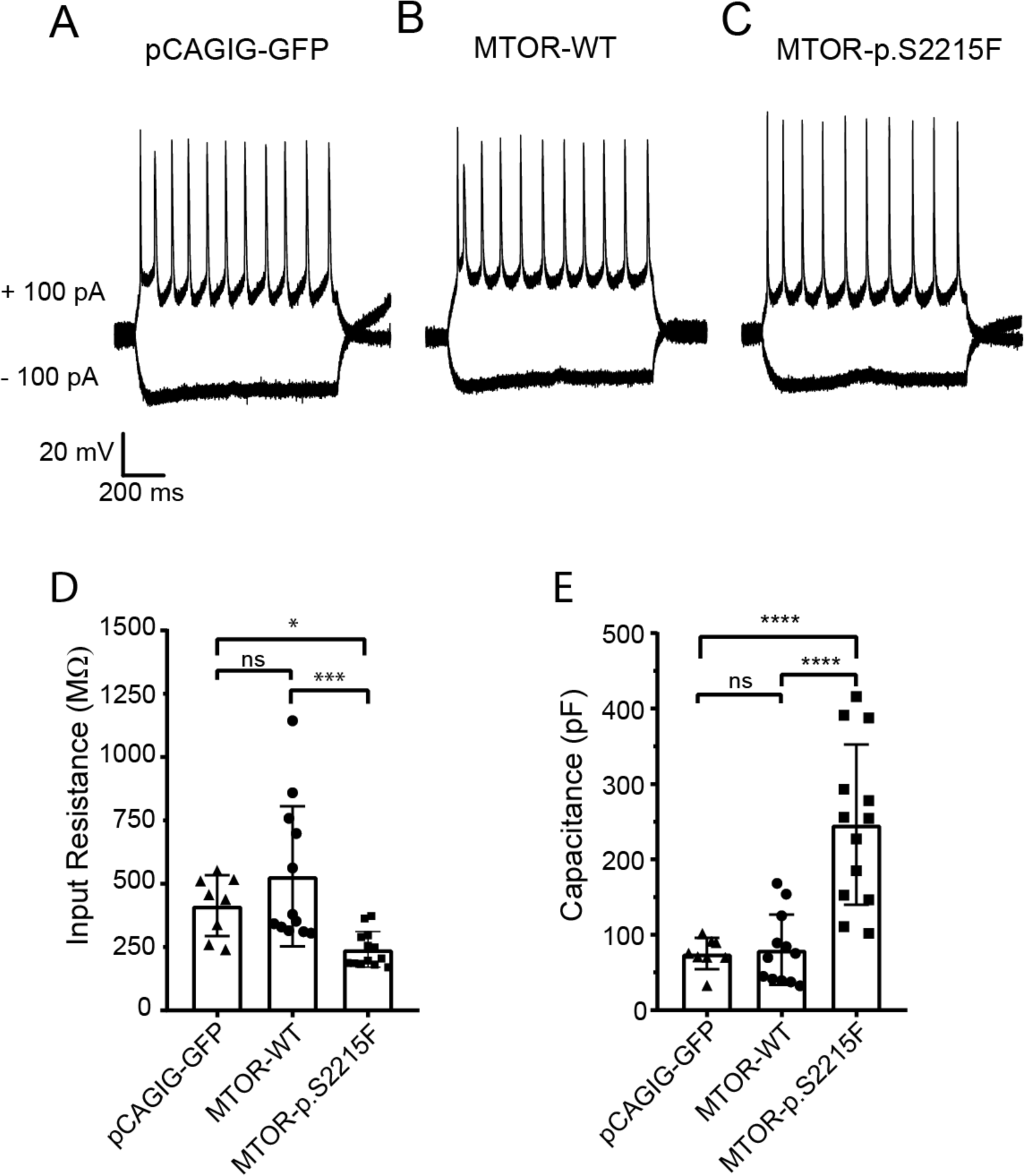
Action potential firing pattern and membrane properties of electroporated neurons. Example traces of action potential firing patterns of spiny stellate neurons recorded in current clamp mode upon depolarizing and hyperpolarizing current injections to the cells and expressing either of (A) GFP only (pCAGIG-GFP), (B) MTOR-WT, and (C) Mutant MTOR-p.S2215F (D) Neuronal input resistance and (E) capacitance values in GFP-only (n=8 neurons from 7 pups), MTOR-WT (n=12 neurons from 11 pups), and mutant MTOR-p.S2215F (n=13 neurons from 11 pups) conditions. ****: p<0.0001; ***: p<0.001; *: p<0.05; ns: not significant. Kruskall-Wallis test followed by Dunn’s post-hoc test.

### Upregulation of NMDAR-mediated currents in mutant MTOR-p.S2215F neurons

Next, we analyzed spontaneous excitatory post-synaptic currents (sEPSCs) at -75 mV (reversal potential of GABA receptor–mediated current) in Mg^2+^-free ACSF to access contribution of AMPA-Receptors (AMPARs) and NMDAR-mediated currents to the total sEPSCs. Mutant MTOR-p.S2215F neurons looked more active than MTOR-WT neurons (Figure 2A) and analysis of the normalized to the peak averaged sEPSCs revealed an increased contribution of the slow component to the total sEPSCs (Figure 2B), most likely indicating elevation of the NMDAR-mediated currents with slow decay kinetics. Importantly, sEPSCs from the non-electroporated, GFP^-^ neurons adjacent to electroporated, GFP^+^ mutant MTOR-p.S2215F neurons were not different from MTOR-WT neurons and from GFP-only neurons (Figure S5). The increased contribution of NMDARs in mutant cells was further confirmed in experiments with APV (50 µM), a selective blocker of total NMDAR-mediated currents. While no change in the amplitude of AMPA receptor (AMPAR)-mediated component of the total sEPSC in the presence of APV was observed (Figure 3A,B), the ratio of the NMDA/AMPA amplitudes significantly increased from 0.3412 ± 0.2751 in MTOR-WT neurons to 1.635 ± 0.5749 in mutant MTOR-p.S2215F neurons (p = 0.0006; Mann-Whitney test, two-tailed) (Figure 3C).

**Figure 2.**
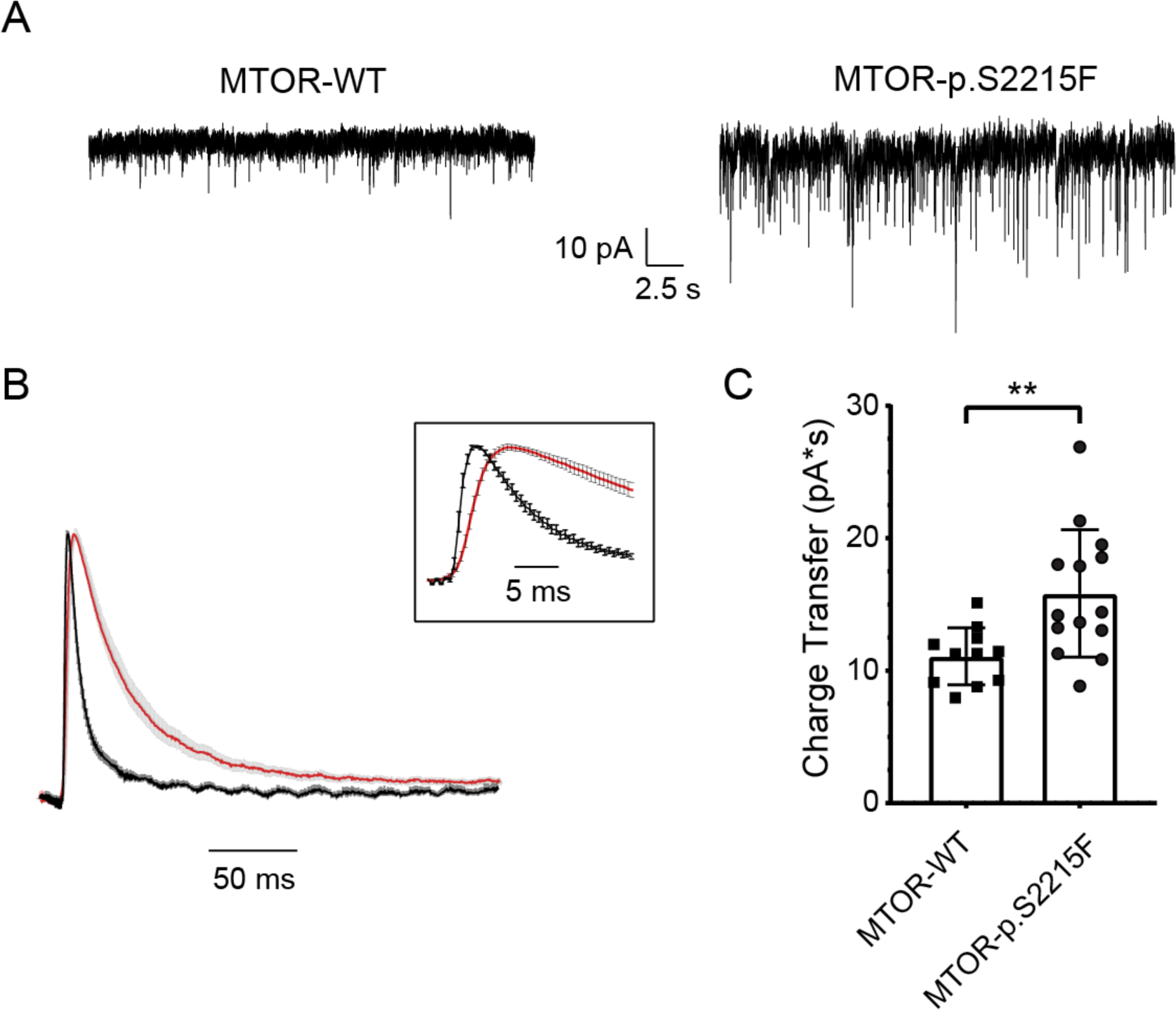
Spontaneous excitatory post-synaptic currents (sEPSCs) in electroporated neurons. (A) Representative traces of sEPSCs in spiny stellate neurons expressing MTOR-WT (left) and mutant MTOR-p.S2215F (right) recorded in Mg^2+^-free ACSF at -75 mV (B) Averaged sEPSCs normalized to the peak for the WT (black) and the mutant (red) neurons. Note a slower current decay and rise times (inset) for mutant neurons indicating increased contribution of NMDAR-mediated component on sEPSC amplitude. Values are given as mean ± SEM (C) Total charge transfer through the sEPSCs normalized to the peak and calculated as an area under the composite sEPSC curve within 200 ms. MTOR-WT neurons, n = 11; mutant MTOR-p.S2215F neurons, n = 14. **: p<0.01. Mann Whitney test, two-tailed.

**Figure 3.**
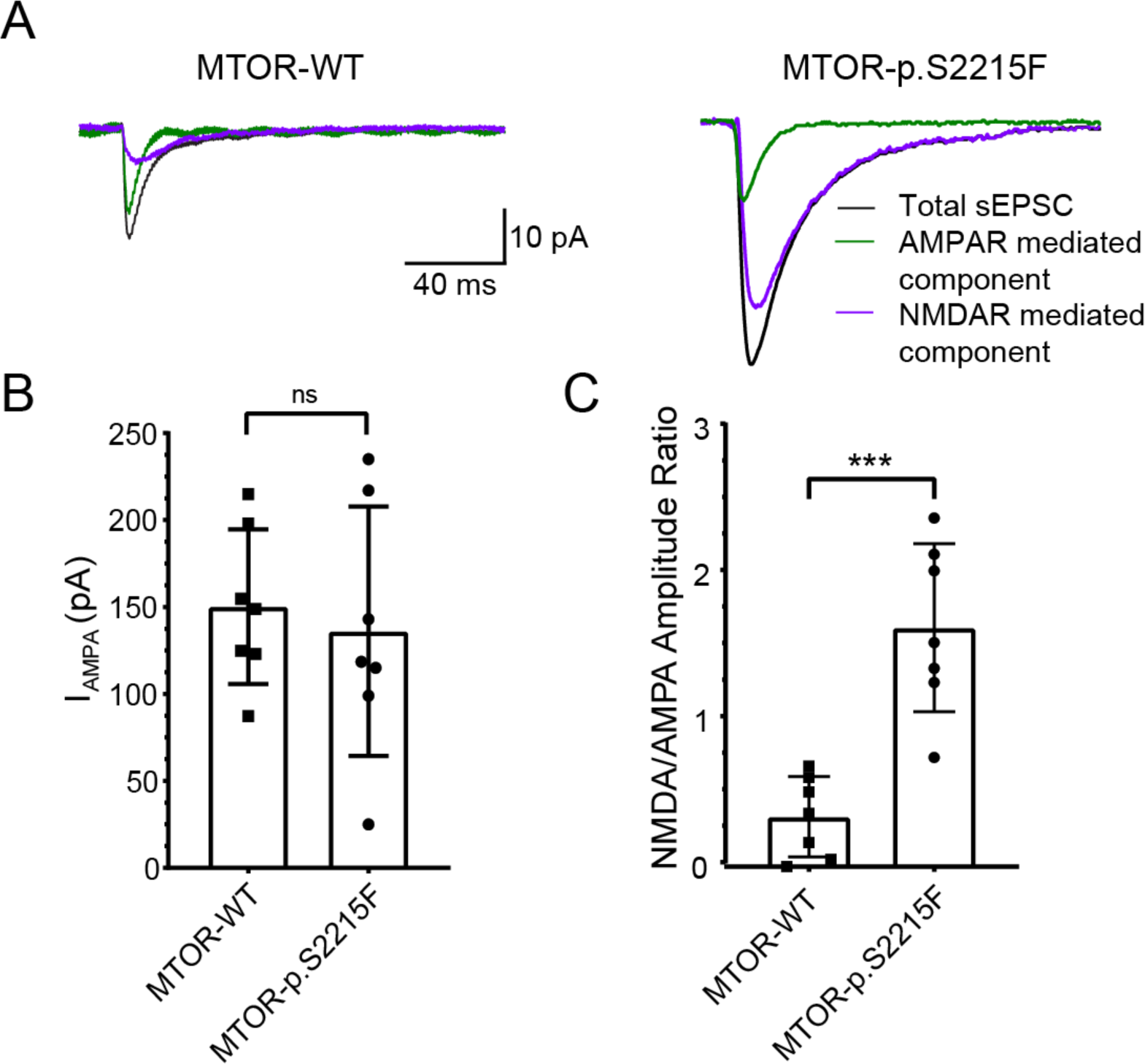
NMDA to AMPA amplitude ratio in sEPSCs in electroporated neurons. (A) Representative traces of averaged sEPSCs recorded in MTOR-WT (left) and p.S2215F-MTOR (right) spiny stellate neurons in: black curve, drug-free (No drug) condition, representing total component of sEPSC; green curve, after adding NMDAR blocker APV (50µM), representing AMPAR-mediated component of sEPSC; and purple curve, after substraction of the AMPAR- mediated component from total sEPSC, representing NMDAR-mediated component of the sEPSC. Note the increased amplitude of the NMDAR-mediated component in the mutant MTOR-p.S2215F neurons while the AMPAR-mediated component remains as in MTOR-WT neurons (B) Summary data for AMPA component amplitude (IAMPA) in MTOR-WT (n = 7) and MTOR-p.S2215F (n = 7 ) neurons. (C) Summary data for NMDA/AMPA amplitude ratios in MTOR-WT (n = 7) and in mutant MTOR-p.S2215F (n = 7) neurons. ***: p<0.001; ns : not significant. Mann Whitney test, two-tailed.

To discriminate between the respective contributions of different NMDAR subunits in increased NMDAR-mediated currents with slow decay kinetics in the total sEPSCs, we also used the following subunit-selective NMDAR blockers: DQP 1105 (10 µM) for GluN2C/D-mediated component, and Ro 25-6981 (1 µM) for GluN2B-mediated component. Comparison of the averaged sEPSC areas revealed that application of DQP 1105 significantly reduced the total sEPSC area while Ro 25-6981 did not have any significant additional effect (No drug vs DQP: p = 0.0054; DQP vs DQP+Ro: p = 0.1221, mixed-effect analysis with Tukey’s multiple comparison test ) (Figure 4). As a possible contribution of GluN2D in the recorded sEPSCs was excluded based on the decay kinetics of the component inhibited by DQP 1105, these results suggested a major contribution of GluN2C- mediated currents to increased NMDAR-mediated component of the total sEPSCs. Also and as expected, daily administration from P8 to P15 of the mTOR inhibitor rapamycin to the pups prior to the patch clamp recordings restored proper neuronal input resistance and membrane capacitance (Figure 5A-C), and led to decreased contribution and faster decay kinetics of NMDAR-mediated currents in mutant neurons (Figure 5D-F); consistent with the lack of GluN2C-mediated effects in this rescue condition, DQP 1105 did not have any significant impact on total sEPSC (Figure S6).

**Figure 4.**
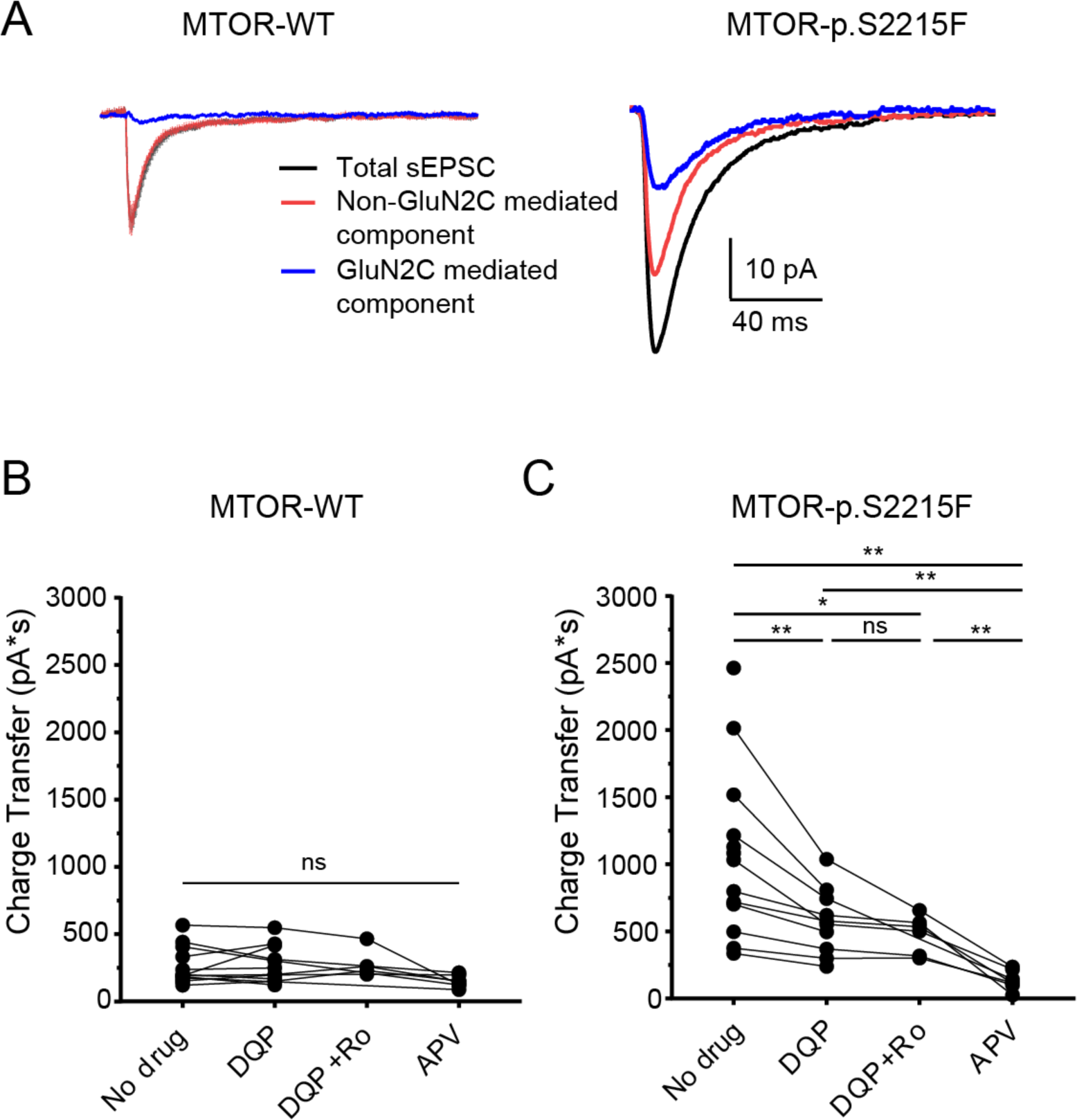
Functional upregulation of GluN2C-containing NMDARs in MTOR p.S2215F electroporated neurons. (A) Representative traces of averaged sEPSCs recorded in MTOR-WT (left) and MTOR-p.S2215F (right) spiny stellate neurons in: black curve, drug-free (No drug) condition, representing total component of sEPSC; red curve, after adding GluN2C-mediated NMDAR blocker DQP 1105 (10 µM), representing the non-GluN2C-mediated component of sEPSC; and blue curve, after subtraction of the non-GluN2C-mediated component from total sEPSC, hence representing GluN2C-mediated component of sEPSC (B,C) Charge transfers were calculated in (B) MTOR-WT neurons and in (C) mutant MTOR-p.S2215F neurons, as areas between peak and 200 ms after the peak of the total sEPSC: with No drug; with GluN2C blocker DQP1105 (DQP; 10 µM); with DQP1105 (DQP; 10 µM) + GluN2B blocker Ro25-6981 (DQP+Ro; 1 µM); and with total NMDAR blocker APV (50 µM). **: p<0.01; *: p<0.05; ns: not significant. Mixed effect analysis with multiple comparison test.

**Figure 5.**
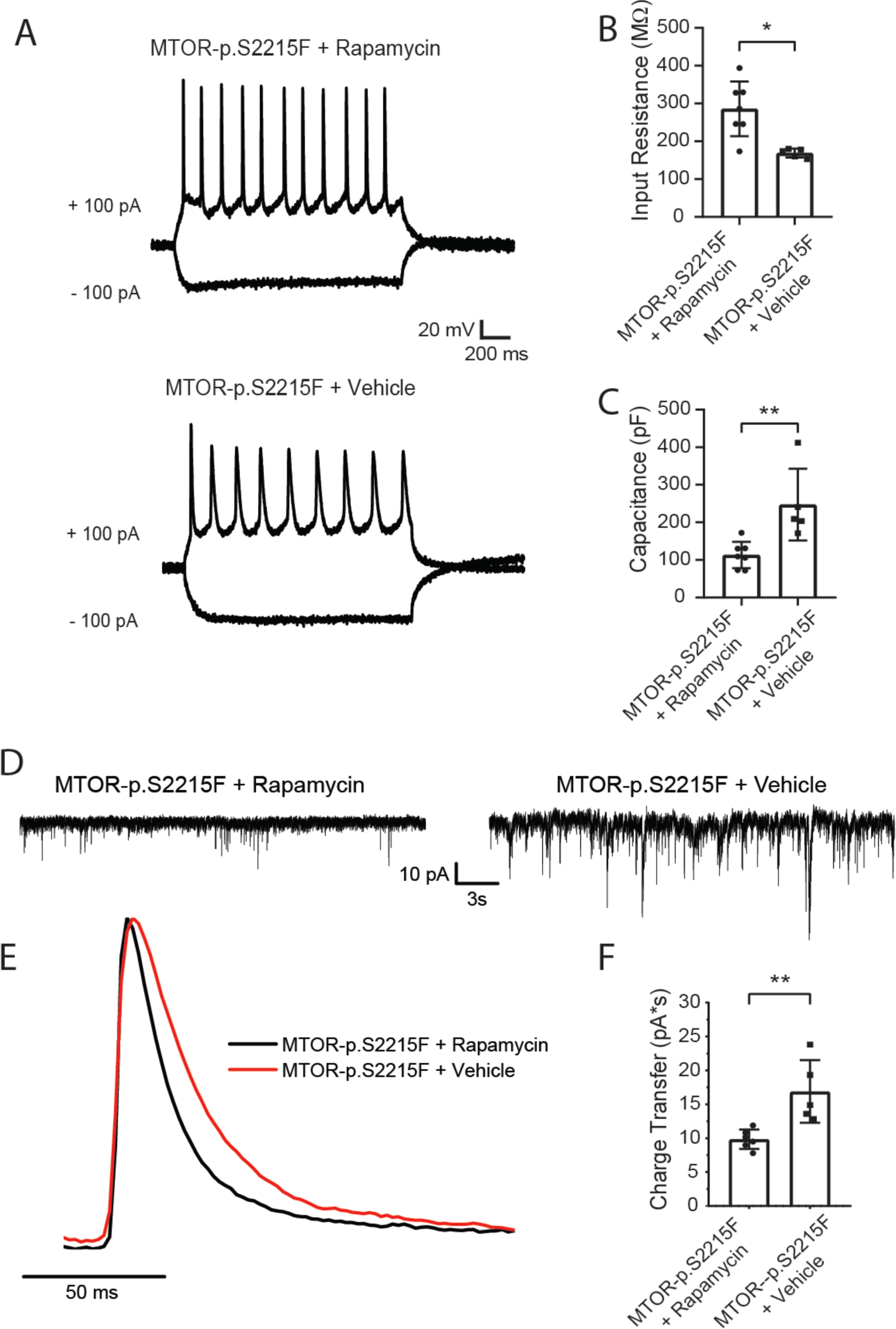
Rapamycin restores intrinsic properties and decreases the charge transfer of normalized sEPSCs in MTOR-p.S2215F neurons. (A) Example traces of action potential firing patterns of spiny stellate neurons expressing MTOR-p.S2215F from pups treated with rapamycin (up) or vehicle (bottom), all recorded in current clamp mode upon depolarizing and hyperpolarizing current injections (B) Neuronal input resistance and (C) capacitance values in MTOR-p.S2215F + Rapamycin (n = 7 neurons from 4 pups) and MTOR-p.S2215F + vehicle (n = 5 neurons from 3 pups) conditions. (D) Representative traces of sEPSCs from rapamycin (left) or vehicle (right) treated neurons expressing MTOR-p.S2215F, recorded in Mg^2+^-free ACSF at -75 mV (E) Averaged sEPSCs normalized to the peak for the rapamycin (black) and vehicle (red) neurons. Note a slower current decay for neurons from rapamycin-treated pups indicating increased contribution of NMDAR–mediated component on sEPSC amplitude (F) Total charge transfer through the sEPSCs normalized to the peak and calculated as an area under the averaged sEPSC curve within 200 ms. MTOR-p.S2215F + Rapamycine, n = 6 neurons from 4 pups; MTOR-p.S2215F + vehicle, n = 5 neurons from 3 pups. **: p<0.01; *: p<0.05. Mann Whitney test, two-tailed.

### MTOR-p.S2215F variation causes spontaneous and time-restricted spiking hyperactivity driven by NMDAR-GluN2C currents

Increased sEPSC area associated with increased contribution of the GluN2C-mediated component with the slow decay kinetics indicated an increased integration time window for excitatory inputs, providing excessed spiking activity of the mutant neurons. Indeed, current clamp experiments showed an increased and longer lasting spiking activity in mutant cells that was reduced upon DQP 1105 application (Figure S7). We thus decided to more precisely address the impact of increased GluN2C-mediated neuronal currents on focal network activity. To this aim, we studied the spontaneous spiking activity of the electroporated area using extracellular recordings in coronal slices from rat pups between P14 and P16. A significant increase of spontaneous spiking activity frequency was observed in MTOR-p.S2215F slices compared to MTOR-WT slices (MTOR-p.S2215F: 0.8480 ± 0.5872 Hz; MTOR-WT: 0.3795 ± 0.5030 Hz; p = 0.0434; Mann-Whitney test, two-tailed) (Figure 6A,B). Importantly, this increased spiking activity was significantly inhibited when slices were acutely treated with DQP 1105 in MTOR-p.S2215F condition, as compared with the untreated condition; no additional effect was observed upon complete block of NMDARs with APV (no drug *vs* DQP: p = 0.0417; no drug *vs* APV: p = 0.7907; Friedman test with multiple comparison) (Figure 6C,D), thus indicating that abnormal network activity in the focal area containing a mosaic of MTOR mutant cells was mostly dependent on GluN2C-mediated NMDARs. In line with their favorable impact on NMDAR-mediated currents in mutant neurons, prior injections of rapamycin to the pups *in vivo* during seven consecutive days prevented against increased spiking activity in MTOR-p.S2215F slices (MTOR-p.S2215F, untreated: 1.901 ± 0.9979 Hz; MTOR-p.S2215F + Rapamycin: 0.5902 ± 0.3601 Hz; p = 0.0337, Mann-Whitney test, two-tailed) (Figure 6E-G).

**Figure 6.**
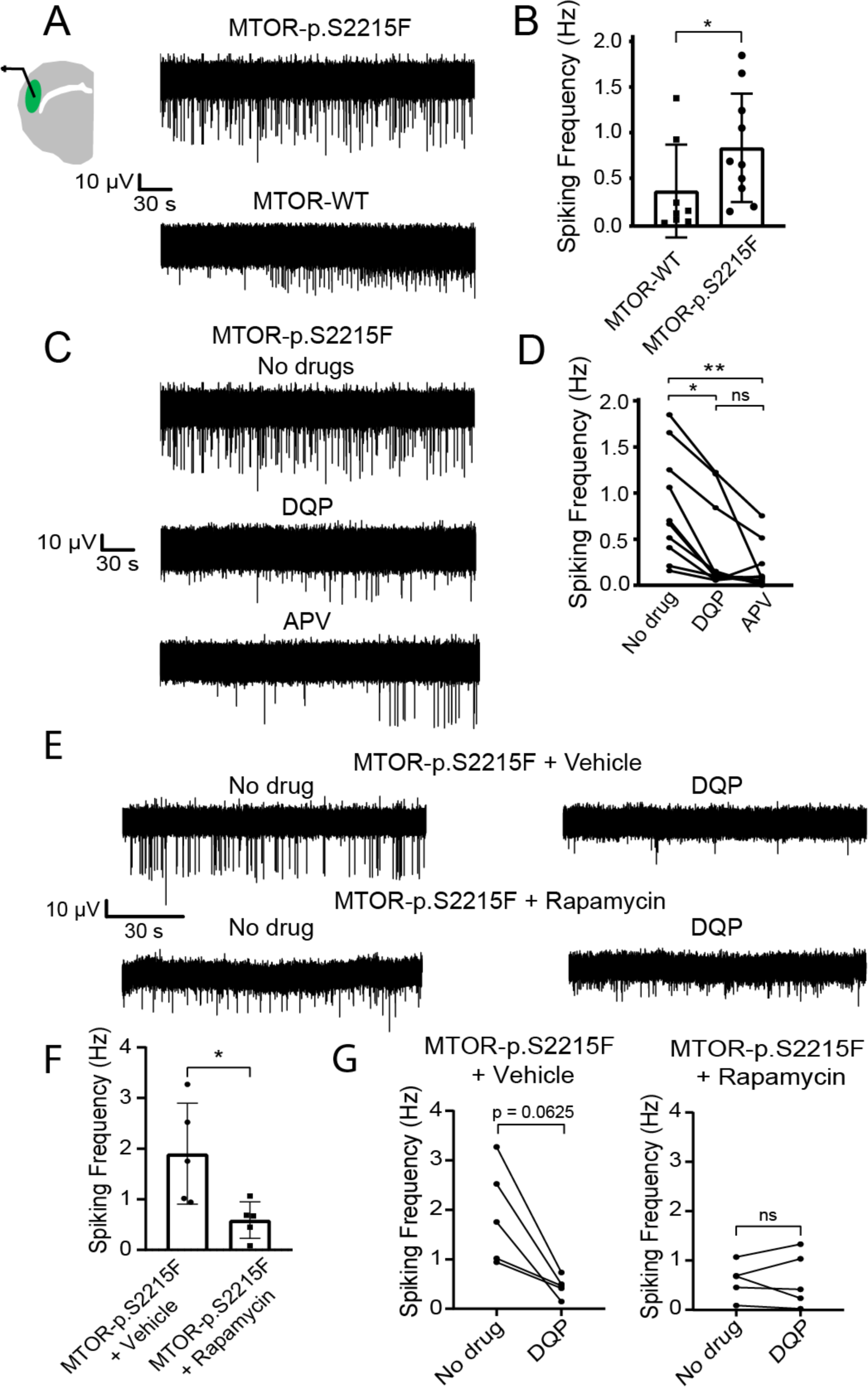
Increased spiking activity in slices from MTOR-p.S2215F electroporated animals. (A) Representative traces of spontaneous activity recorded from green fluorescent electroporated areas in Mg^2+^ containing ACSF medium from (top trace) mutant MTOR-p.S2215F slice and (bottom trace) MTOR-WT slice (B) Summary data for spiking frequency. MTOR-WT, n = 8 slices from 7 pups; MTOR-p.S2215F, n = 10 slices from 9 pups. *: p<0.05. Mann Whitney test, two-tailed (C) Representative traces of spontaneous activity recorded in mutant MTOR-p.S2215F slices (n = 10 from 9 pups) obtained in Mg^2+^ containing ACSF with: (top) no drug; (middle) GluN2C blocker DQP1105 (DQP, 10 μM); and (bottom) total NMDAR blocker APV (50 μM) (D) Summary data for spiking frequency. **: p<0.01; *: p<0.05. Friedman test with Dunn’s multiple comparison test (E) Representative traces of spontaneous activity recorded from the electroporated areas in Mg^2+^ containing ACSF medium from: (top trace) slice from vehicle-treated, mutant MTOR-p.S2215F pup; and (bottom trace) slice from rapamycin-treated, mutant MTOR-p.S2215F pup without (No drug, left) or with DQP 1105 (10 µM, right) (F) Summary data for spiking frequency; n = 5 slices from 2 pups in each condition. *: p < 0,05. Mann-Whitney test, two-tailed (G) Summary data for spiking frequency without or with DQP 1105 for MTOR-p.S2215F + vehicle (left, n = 5 slices from 2 pups) and MTOR-p.S2215F + Rapamycine (right, n = 5 slices from 2 pups). Note the statistical trend towards significant effect of DQP 1105 in the MTOR-p.S2215F + vehicle condition, as expected. Wilcoxon test, two-tailed.

We then decided to examine whether the GluN2C-dependent spontaneous spiking hyperactivity as detected in the area of electroporated neurons in the mutant condition at P14-P16 would be restricted to a critical time-window, as previously reported in *Tsc*^+/-^ heterozygous mice (Lozovaya et al. 2014). Extracellular slice recordings done at P9 did not reveal any significant increase in spiking activity in the electroporated area in the mutant condition (Figure 7A,B); however, such an activity was driven by GluN2C-containing NMDARs, as shown by the significant impact of DQP 1105 on the frequency of action potentials (Figure 7C). Consistently, sEPSCs of MTOR-p.S2215F electroporated neurons at P9 showed a strong DQP 1105 sensitive component with a deactivation kinetic consistent with GluN2C-containing NMDAR-mediated currents (Figure S8).

**Figure 7.**
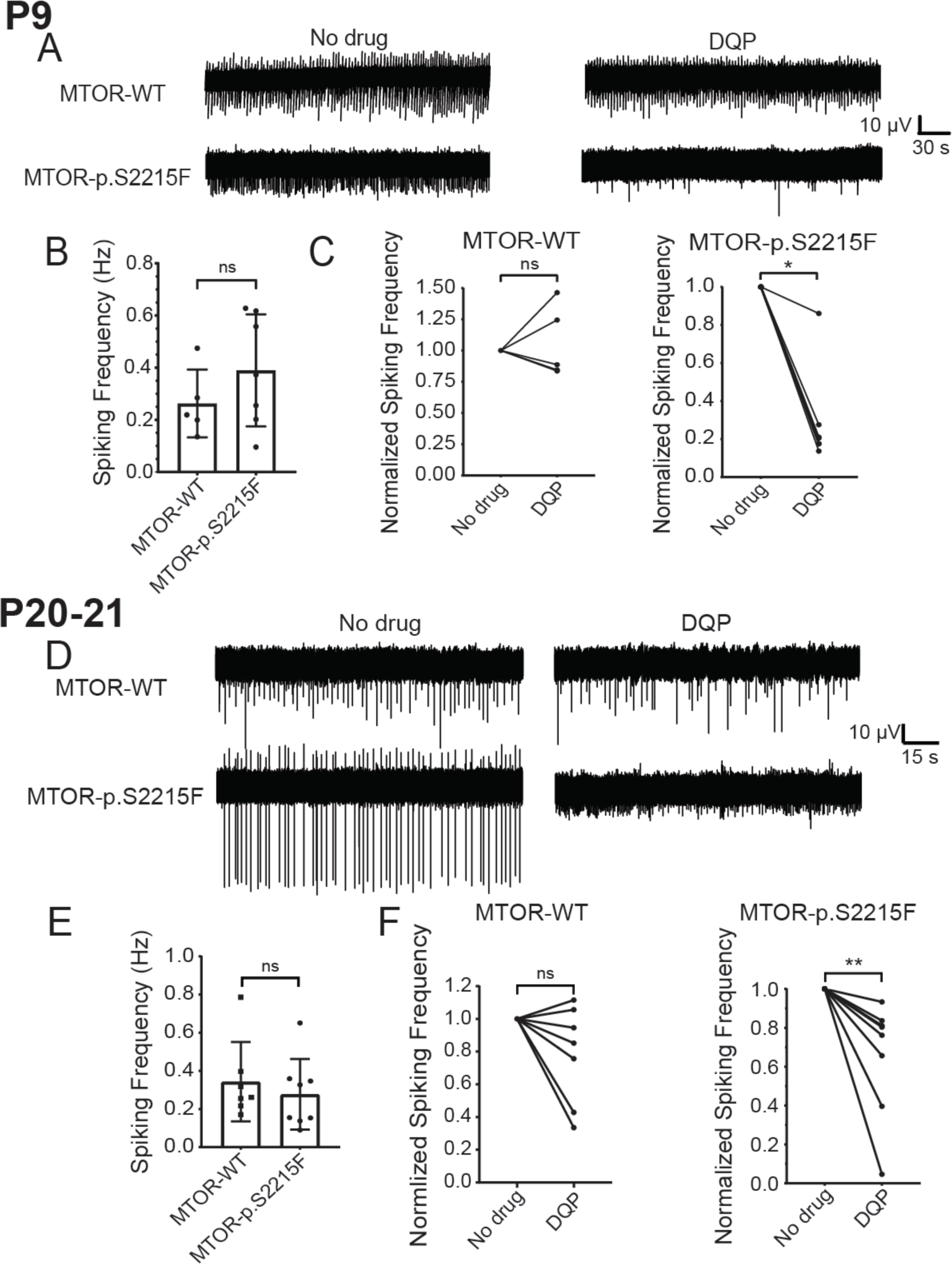
Developmental evolution of GluN2C-dependent increased spiking activity. (A) Representative traces of spontaneous activity recorded from electroporated areas in Mg2+ containing ACSF medium from (top trace) MTOR-WT slice and (bottom trace) MTOR-p.S2215F slice from P9 pups, without (No drug, left) or with DQP 1105 (10 µM, right) (B) Summary data for spiking frequency. MTOR-WT, n = 5 slices from 4 pups; MTOR-p.S2215F, n = 7 from 6 pups. ns: not significant; Mann-Whitney test, two-tailed (C) Summary data for spiking frequency without or with DQP 1105 for MTOR-WT (left) and MTOR-p.S2215F (right). Data shown are values normalized to the No drug condition. *: p<0.05; ns: not significant; Wilcoxon test, two-tailed (D) Representative traces of spontaneous activity recorded from electroporated areas in Mg2+ containing ACSF medium from (top trace) MTOR-WT slice and (bottom trace) MTOR-p.S2215F slice from P20-21 pups without (No drug, left) or with DQP 1105 (10 µM, right) (E) Summary data for spiking frequency. MTOR-WT, n = 7 slices from 4 pups; MTOR-p.S2215F, n = 8 slices from 5 pups. ns: not significant. Mann Whitney test, two-tailed (F) Summary data for spiking frequency without or with DQP 1105 for MTOR-WT (left) and MTOR-p.S2215F (right). Data shown are values normalized to the No drug condition. **: p<0.01; ns: not significant. Wilcoxon test, two-tailed.

As at P9, at P20-P21 spontaneous spiking activity of slices from mTOR-p.S2215F electroporated animals, even though not increased as compared with MTOR-WT slices (Figure 7D,E), was qualitatively modified, as it showed sensitivity to DQP 1105, while activity in MTOR-WT slices was not modified by DQP 1105 (Figure 7F).

## Discussion

Genetic defects leading to excessive activation of the mTOR pathway represent a well-established and important cause of various types of epileptic disorders, notably in the context of brain malformations (Aronica et al., 2012; Baulac, 2016; Blümcke et al., 2021; Crino, 2015).

However, the relationship with epileptogenesis and with the underlying hyperexcitability of neuronal networks remains to be clarified (Nguyen and Bordey, 2021; Represa, 2019). In the recent years, the involvement of somatic mTOR variants in FCD type II has been increasingly recognized (Blumcke et al., 2021a). We previously reported that the mosaic expression of a somatic, recurrent p.S2215F MTOR variant in neurons of the rat neocortex was associated with neuronal dyslamination and dysmorphia, two key hallmarks of FCD type II, and with increased S6 phosphorylation taken as a readout of mTORC1 overactivation (Pelorosso et al., 2019). In this model, we now show that, in rat pups aged P14-P16, spiny stellate neurons expressing the mutant MTOR protein displayed intrinsic abnormalities and an excessive activation of NMDAR-mediated component of total sEPSCs not observed in MTOR-WT neurons, and sustained by GluN2C-containing NMDARs. Such an aberrant activity resulted in increased spontaneous spiking activity *in vitro*, revealing excessive local network activity that was restricted to a critical time window between P9 and P20. Slow decay kinetics of NMDAR GluN2C-mediated currents, leading to an increased synaptic integration, would facilitate neuronal network synchronicity. Hence, the GluN2C-dependent hyperexcitability previously reported in TSC (Lozovaya et al., 2014), which is a mTORopathy caused by constitutional loss-of-function defect in either of *TSC1* or *TSC2* genes, is now expanded to another type of mTORopathy and to another genetic context, namely FCD type II caused by a somatic, gain-of-function MTOR variant.

### From cell autonomous GluN2C-related anomalies to local hyperexcitability

Studies in different models of mTORopathies have pointed for nearly as many possible mechanisms of neuronal and network hyperexcitability (Nguyen and Bordey, 2021). In our model, no change was detected in non-electroporated neurons adjacent to the dysmorphic, electroporated neurons expressing mutant MTOR. This is in line with previous findings in other mosaic models of mTOR-related brain disorders, where cell autonomous mechanisms were also reported (Hsieh et al., 2016; Park et al., 2018; Wu et al., 2022). We do not exclude the possible participation of more distant cells or even of normotopic cortex to FCD-related hyperexcitability; indeed, non cell autonomous mechanisms were previously reported in other models of brain cortical malformations, including mTOR-related ones (Kim et al., 2019; Koh et al., 2021). In FCD tissues from patients and in animal models, the altered expression of GABA receptors (Calcagnotto et al., 2005; Crino et al., 2001), the increased expression of Kv1.1 potassium channels (Nguyen and Anderson, 2018) or the ectopic expression of HCN4-mediated Ih currents in mutant neurons (Hsieh et al., 2016) have also been proposed. When different genetic models of focal MTORopathies were recently compared, both shared (e.g. aberrant HCN4 channel expression, increased sEPSCs) and divergent (e.g. different characteristics of the recorded sEPSCs) findings on the electrophysiological properties of the mutant neurons were found (Nguyen et al. 2023); of note, the brain area (medial prefrontal cortex), the neuronal type (pyramidal neurons), and the developmental timepoint of the recordings (P28-P43) were all different from those studied here.

Our data point for a cell-autonomous role of overexpressed GluN2C-mediated NMDAR currents in spiny stellate neurons. Spiny stellate neurons are considered as the main acceptors of thalamic sensory inputs in the somatosensory cortex (Feldmeyer, 2012; Feldmeyer et al., 1999). They are excitatory cells that are highly interconnected within layer 4 and also have outputs in all cortical layers in their own barrel but also in adjacent ones. Considering the connectivity of spiny stellate neurons and the specific properties of GluN2C-gated NMDAR currents, the overexpression of GluN2C-containing NMDAR currents in spiny stellate neurons is very likely responsible for an increased probability of synchronising neuronal activities in this FCD model. Noteworthingly, increased GluN2C-mediated NMDAR currents were also recorded in spiny stellate neurons in *Tsc*^+/-^ mice (Lozovaya et al., 2014). Of note, also, GluN2D-containing NMDA receptors would also be sensitive to DQP 1105 and to other currently used GluN2C/D inhibitors (Acker et al., 2011;Lozovaya et al., 2014; Nouhi et al., 2018; Wang et al., 2020); however, the participation of GluN2D-NMDARs was excluded as the decay kinetics of the component inhibited by DQP 1105 corresponded to GluN2C and not GluN2D contribution (Paoletti et al., 2013; Seeburg et al., 1995).

### From mTORopathies to hyperexcitability: GluN2C-mediated neuronal NMDAR currents as a unifying mechanism

We had previously reported on the pathogenic role of GluN2C-mediated NMDARs in hyperexcitability of a TSC mouse model and of resected tissues from TSC patients (Lozovaya et al., 2014). Overexpression of *GRIN2C*, which encodes the GluN2C subunit, and GluN2C-dependent hyperexcitability had already been detected in FCD tissues, albeit of undetermined genetic status (Crino et al., 2001; Lozovaya et al., 2014). Here, we now demonstrate that GluN2C drives focal hyperexcitability in the context of a MTOR pathogenic variant that is recurrently detected in FCD type II patients. Altogether and as mTOR variants represent the majority of known somatic genetic causes of FCD type II, the GluN2C-related pathomechanism of focal mTORopathy as revealed here, might well be shared in common by most FCDs type II. Noteworthingly, treatment of the pups with the mTOR inhibitor rapamycin prior to the recordings prevented against hyperactive NMDARs and GluN2C functional upregulation, and against increased spiking activity. This confirms that the p.S2215F variant does not impact on MTOR sensitivity to rapamycin, as previously reported (Xu et al., 2016). MTOR is a member of both the mTORC1 and the mTORC2 complexes, and it was shown that rapamycin exerts its inhibitory effect on mTOR activity by targeting the MTOR protein in the mTORC1, but not in the mTORC2 complex (Seto, 2012). Hence, our data indicate an important contribution of the mTORC1 pathway to pathogenesis in our model. Moreover, our data point on postsynaptic upregulation of NMDAR-mediated currents, which is consistent with previous findings showing that mTORC1 acted on glutamatergic synaptic transmission *via* a postsynaptic mechanism whereas mTORC2 affected presynaptic parameters (McCabe et al., 2020). Still, our data do not exclude the participation of mTORC1-independent pathological processes. For instance, it was shown that inhibition of mTORC2 could reduce the neurobehavioral abnormalities of *Pten*-deficient mice (Chen et al., 2019) and, even more interestingly, of adult mice harboring the p.S2215F MTOR variant in a similar mosaic configuration as in the present study (Okoh et al., 2023), suggesting that both mTORC1- and mTORC2-related pathomechanisms would be operating, possibly at different developmental timepoints, in FCD caused by the p.S2215F MTOR variant.

Nevertheless, as mTORC1 upregulation ultimately converges on dysregulated phosphorylation of shared downstream products (e.g. RPS6, 4EBP1), the GluN2C-mediated hyperexcitability of neuronal networks might expand to an even broader spectrum of mTORC1- related disorders than TSC and FCD type II. Whereas this obviously warrants expanded confirmation in the corresponding models of mTORopathies, this would make it even more crucial to identify in the future how mTOR hyperactivation leads to increased GluN2C-mediated NMDAR currents.

While we and others have previously demonstrated increased expression of the corresponding *GRIN2C* gene in human resected tissues and in mice (Crino et al., 2001; Lozovaya et al., 2014), and while the role of the mTOR pathway in transcriptional regulation is well recognized (Martina et al., 2012; Roczniak-Ferguson et al., 2012; Zhang et al., 2017), what transcription and other (e.g. chromatin-regulating proteins) factors would sustain *GRIN2C* dysregulation should be questioned in the future. Furthermore, post-transcriptional effects of mTOR dysregulation on the expression of GluN2C-mediated currents, such as increased mTOR-dependent protein translation or membrane trafficking, may be involved as well. Interestingly, the spiny stellate neurons that had been electroporated *in utero* to express the wild-type isoform of MTOR did not show any quantitative or qualitative anomaly in NMDAR-mediated currents, as compared with the GFP-alone condition.

GluN2C-mediated currents have been detected by single channel recordings in spiny stellate neurons of the somatosensory cortex in naive mouse pups (Binshtok et al., 2006; Scheppach, 2016); hence, mTOR upregulation caused by MTOR-p.S2215F variant would be needed to trigger overexpression of GluN2C-mediated currents detectable by whole cell recordings. This is in line with the lack of dyslamination and of neuronal dysmorphism as observed here upon expression of wild-type MTOR, and with previous findings showing that overexpression of wild-type MTOR *per se* did not influence neuronal migration and morphology (Kassai et al., 2014; Kim et al., 2019; Lim et al., 2015; Pelorosso et al., 2019).

### A critical time-window for GluN2C-related pathomechanisms of mTORopathies?

As for epileptic activity in *Tsc1*^+/-^ mice (Lozovaya et al., 2014), local hyperexcitability as recorded at P14-P16 looked transient as it was neither detected at earlier (at P9) nor at later (P20- P21) stages. Hence, GluN2C-mediated hyperactivity would follow the same developmental pattern in TSC and in FCD type II, being restricted to a similar critical developmental period. Interestingly, even though GluN2C-mediated activity was already upregulated and significantly contributed to NMDAR-mediated currents in spiny stellate neurons at P9, this did not induce local network hyperactivity at that early stage. Also, compensatory mechanisms possibly driven by developmental changes in the relative expression patterns of other NMDARs subunits, with GluN2A becoming predominant over GluN2B with age (Liu et al., 2004; Monyer et al., 1994) and conferring faster kinetics to NMDARs, might explain why hyperexcitability was no longer detected at the end of the third week of life.

The data obtained here make it the GluN2C-containing NMDAR a possible target in FCD type II caused by somatic variants of the mTOR pathway, as also previously suggested for autosomal dominant TSC (Gataullina et al., 2022; Lozovaya et al., 2014). The importance of GluN2C in the pathogenesis of FCD might even expand beyond the spectrum of mTORopathies, as very recently suggested by the detection of a likely pathogenic, somatic *GRIN2C* variant in a patient with FCD type I (Chung et al., 2023). Our results also indicated that the GluN2C anomalies observed in our model of FCD followed the same developmental pattern as for *Tsc*^+/-^ mice, suggesting the existence of a critical time window for GluN2C-mediated pathomechanisms and for related rescue strategies.

While efforts towards the development of novel GluN2C selective inhibitors are ongoing (Gataullina et al. 2022; D’Erasmo et al. 2023), possible off-target effects of GluN2C inhibition might also be considered. While not detectable at birth, mouse GluN2C expression is enriched in the second postnatal week in the cerebellum, the thalamic reticular nucleus and the olfactory bulb (Monyer et al., 1994; Ravikrishnan et al., 2018). Systemic administration of GluN2C inhibitors would likely impact on NMDARs expressed in such distant brain areas. Indeed, GluN2C influences cortical excitation/inhibition balance and neuronal oscillations (Gupta et al., 2016). Local administration at the site of the lesion would likely prevent against such broad side effects; however, GluN2C- mediated currents are also detectable in spiny stellate neurons in physiological conditions, as shown by single channel recordings (Binshtok et al., 2006), and can also be expressed in non-neuronal cells such as astrocytes and microglia (Alsaad et al., 2019; Lee et al., 2010; Ravikrishnan et al., 2018).

Despite these limitations, our data suggest that early, focal inhibition of GluN2C-mediated currents during a critical time period might be considered in FCD type II (Moloney et al., 2021). As a matter of fact, early interventions against mTOR-related dysregulations are also being considered to prevent against the progression to severe epilepsy in TSC (Aronica et al. 2023).

## Conclusion

Slow decay kinetics of GluN2C-mediated NMDAR currents, by leading to increased synaptic integration, facilitate neuronal network synchronicity. We show here that GluN2C-containing NMDARs drive time-restricted focal hyperexcitability in the context of a MTOR somatic variant that was recurrently detected in FCD type II patients. As somatic variants of the mTOR pathway represent an important cause of FCD type II, the GluN2C-related pathomechanism of focal mTORopathy as revealed here might well be shared in common by most FCDs type II, and beyond in the context of other mTORopathies.

## Author contributions

LP performed most experiments, did most data analyzes of recordings with NB, and wrote the manuscript with PS and NB. EB performed the *in utero* intracerebroventricular electroporations and the EEG recordings. ST participated in IHC experiments and in extracellular electrophysiological recordings and analyses. SB participated in follow-up of the study and in IHC experiments. VC analyzed EEG traces with LP. FW, CC and AR participated in the design and follow-up of the study. PS and NB co-decided on the overall strategy, co-directed the follow-up of experiments and wrote the manuscript. All authors contributed the final version of the manuscript.

## Acknowledgments

We thank E Bouilloc, P Moudery and S Corby for their help at INMED (Mediterranean Institute of Neurobiology) animal core facilities, A Montheil and E Pallesi-Pocachard at INMED Molecular and Cell Biology platform, and F Michel at INMED InMagic platform. We also thank L Aniksztejn for his advices in electrophysiological recordings. L Pineau has been a recipient of a Ministry of Research Aix-Marseille University Doctoral School (ED62) PhD fellowship. S Tarhini is a recipient of an Aix-Marseille University/Amidex/CMA CGM PhD fellowship. This work was supported by INSERM (Institut National de la Santé et de la Recherche Médicale) and by the European Union Seventh Framework Program FP7/2007-2013 under the project DESIRE (grant agreement n°602531).

## Supplementary Material

Table S1

Figures S1-S8

