## Supplementary Figures for "Pathogenic MTOR somatic variant causing focal cortical dysplasia drives hyperexcitability *via* overactivation of neuronal GluN2C NMDA receptors"

**to**

**Figure S1. MTOR-p.S2215F variant leads to spontaneous epileptic episodes in two months old rats.**

(A) Representative traces of EEG recordings from MTOR-WT (top) and MTOR-p.S2215F (bottom) electroporated rats aged two months, showing recurrent spike and wave complexes (magnified on the right side) in MTOR-p.S2215F recordings and occasional single spike and wave (magnified on the right side) in MTOR-WT recordings (B) Top: representative trace of a spontaneous epileptic episode in MTOR-p.S2215F electroporated rat aged two months. Bottom: magnified views of the pre-ictal (left), ictal (middle) and post-ictal (right) phases.

Spike-wave complexes (SWs) were detected more frequently, albeit with great inter-individual variability, in the three MTOR-p.S2215F rats (57.84 ± 55.57 events/h), while fewer SWs (0.5268 ± 0.3762 events/h) were seen in the three MTOR-WT rats. The two mutant MTOR-p.S2215F rats with the more numerous SWs had two epileptic episodes in the 72h of recordings, while the third mutant rat with much less SWs displayed one epileptic episode. The episodes were characterized by a mean duration of 24.75 ± 8.098 s. In contrast ($p = 0.05$, Fisher's exact test), no epileptic episode was ever detected in the three MTOR-WT rats.

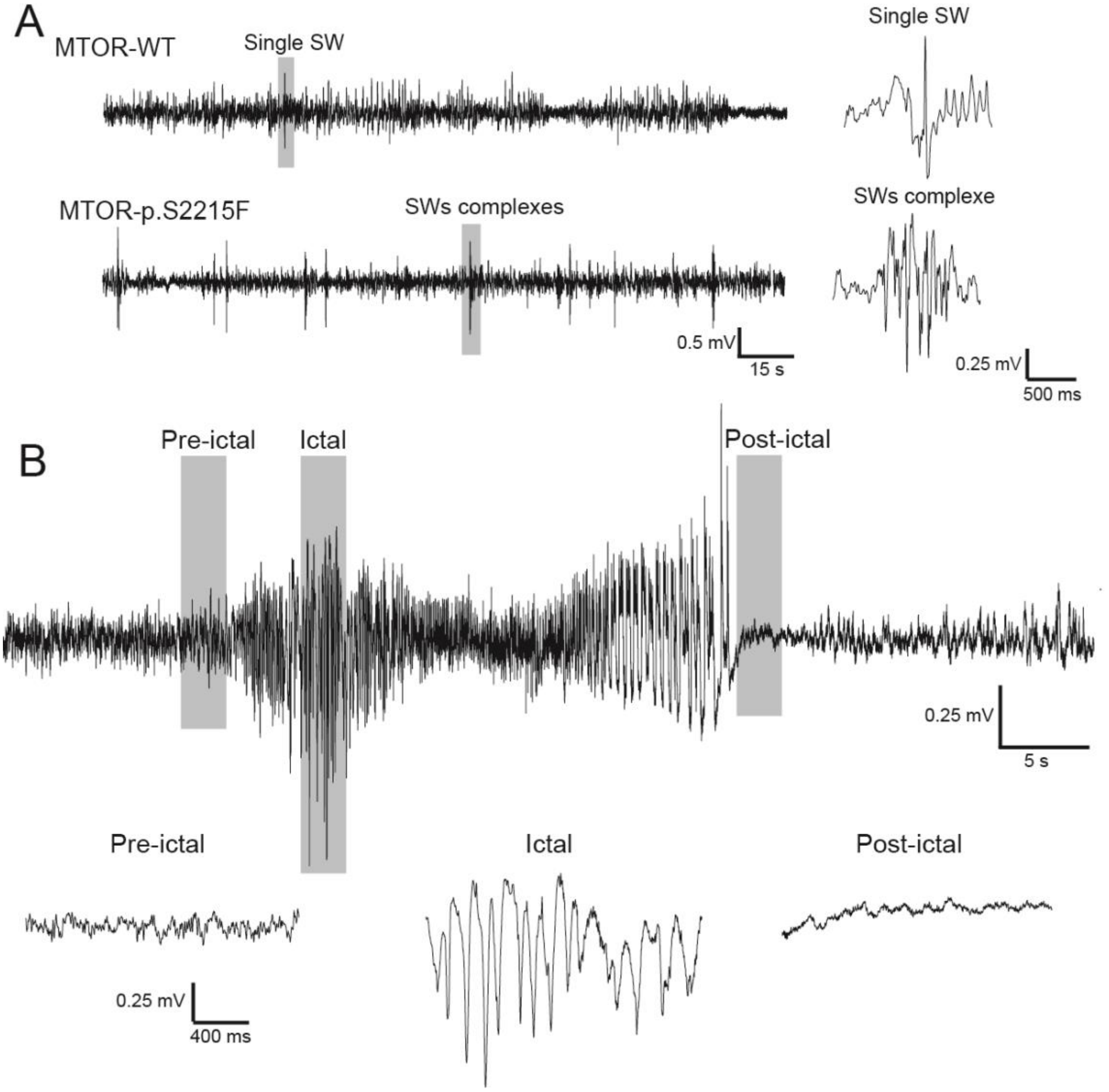

**Material and Method related to Figure S1.**

To implant recording electrodes, rats were anaesthetized with 4.5% isoflurane and maintained at 2.5% isoflurane, placed in stereotaxic frame and prepared for aseptic surgery. A first hole was drilled in the skull at the approximate coordinates of the electroporated primary somatosensory cortex. A screw connected to EEG transmitter was placed in the skull, and reference electrode was placed rostrally in the cerebellum. The screws were attached to the skull with dental cement and scalp closed. After surgery, rats were kept warm and received saline (0.9% NaCl, subcutaneous injection, s.c.), and analgesic buprenorphine (0.03 mg/kg, s.c.) was administrated to minimize animal pain after surgery. After a five days recovery, EEG (amplified (1000×), filtered at 0.16–97 Hz pass, acquired at 500 Hz) was monitored using a telemetric system (Data Sciences International, St. Paul, MN) for three consecutive days (72 h.); food and water were given ad libitum. Recordings were visualized offline using NeuroScore (Data Sciences International, St. Paul, MN). After the recordings, rats were intracardiacally perfused, brains were dissected and post-fixed during 48h. Presence of GFP to assess the success of the electroporation was detected by immunostaining as described below. Detection of spike-wave complexes (SWs) and of seizures was performed in 72-hours recordings using Neuroscore® software (Data Sciences International, St. Paul, MN) followed by visual inspection to remove false-positive events. Detection of SWs was done using the dynamic threshold spike detection (threshold ratio: 2; maximum threshold: 20; minimum value: 300 µV) and spike train detection feature (minimum spike interval: 0.05s; maximum spike interval: 0.15s; minimum train duration: 0.5s; train join interval: 1s; minimum number of spikes: 4). Seizure detection was performed by eye using the periodogram feature (10s epoch). Events presenting spike trains, a frequency higher than 35 Hz and duration of more than 6s were visually inspected by two independent experimentators and identified as seizures.

**Figure S2. MTOR-p.S2215F electroporated neurons display increased soma size and dendritic tree compared to MTOR-WT electroporated neurons.**

(A) Morphological reconstruction of the soma (red) and dendritic tree (black) of biocytin-filled MTOR-WT (left) and MTOR-p.S2215F (right) electroporated (GFP+) neurons from P14-P16 pups. Quantification of (B) soma volume (C) dendrites number and (D) dendrites surface area. MTOR-WT, n = 8 neurons from 7 pups; MTOR-p.S2215F,  n = 8 neurons from 8 pups. **: $p < 0.01$; *: $p < 0.05$. Mann-Whitney test, two-tailed.

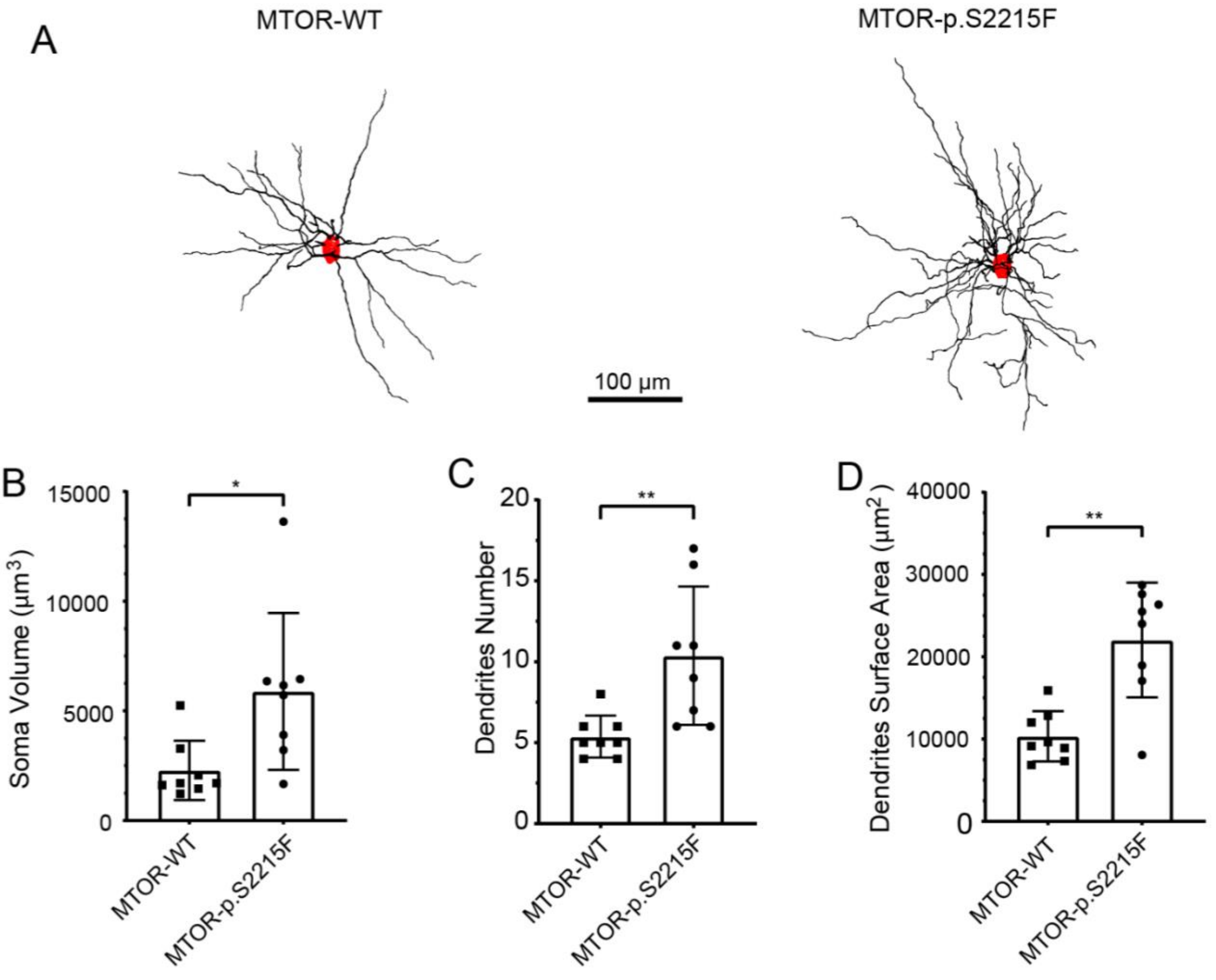

**Figure S3. Summary of intrinsic properties investigated in pCAGIG-GFP, MTOR-WT and MTOR-p.S2215F electroporated neurons.**

Quantification of (A) first action potential (AP) amplitude, (B) first AP halfwidth, (C) AP threshold, (D) rheobase and (E) resting membrane potential. MTOR-WT, n = 12 from 11 pups; pCAGIG-GFP, n = 8 from 7 pups; MTOR-p.S2215F, n = 13 from 11 pups. No significant difference was found between the three conditions. Kruskall-Wallis test with Dunn's multiple comparison test.

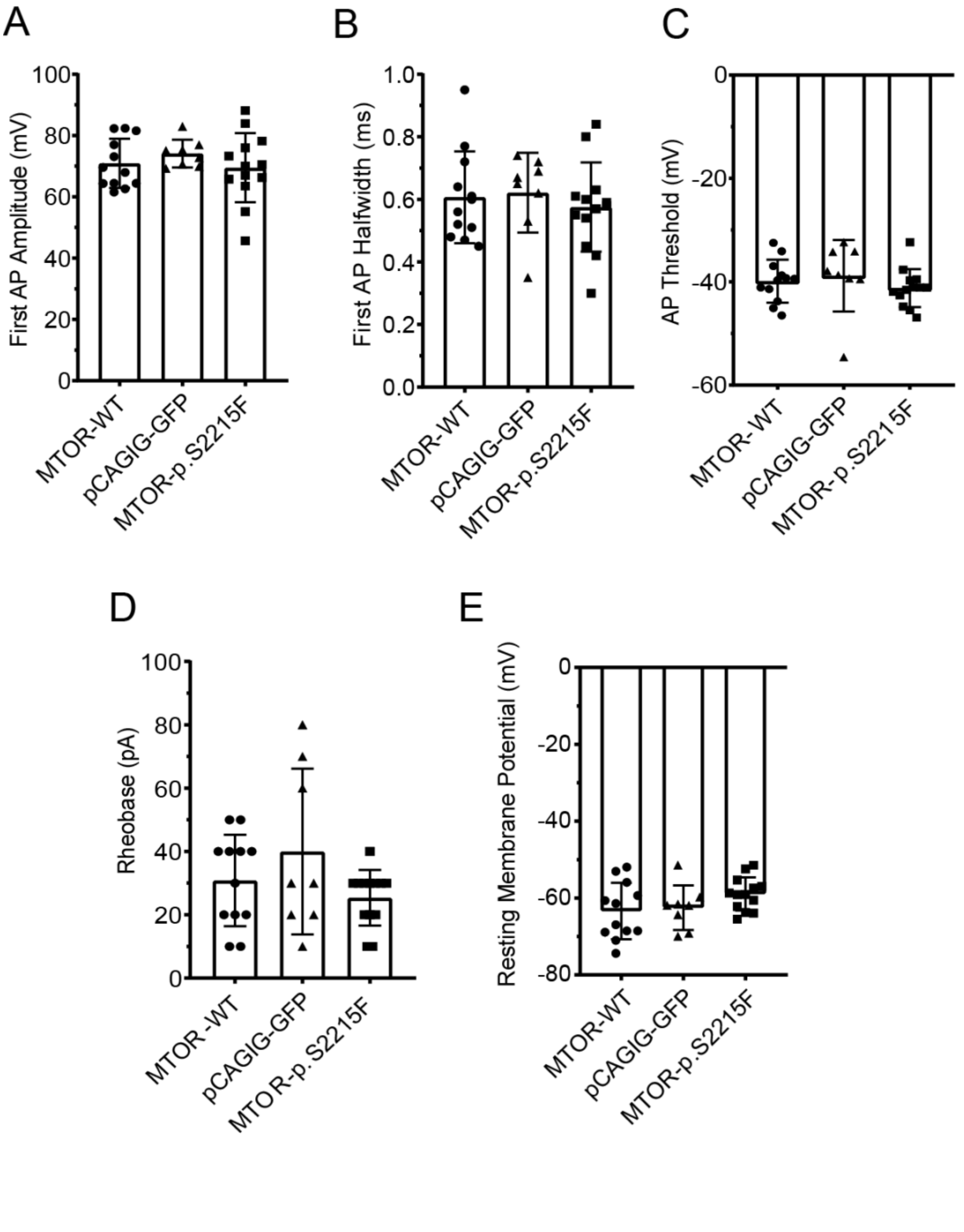

**Figure S4. Summary of intrinsic properties of MTOR-p.S2215F electroporated (GFP+) and adjacent, non-electroporated (GFP-) neurons.**

Quantification of the (A) first action potential (AP) halfwidth, (B) capacitance, (C) first AP amplitude, (D) input resistance, (E) rheobase, (F) resting membrane potential, (G) AP threshold. MTOR-p.S2215F GFP+, n = 13 from 11 pups; non-electroporated (GFP-) neurons adjacent to MTOR-p.S2215F GFP+ neurons, n = 20 from 16 pups; MTOR-WT GFP+, n = 12 from 11 pups. ****, $p < 0.0001$; **, $p < 0.01$. Kruskall-Wallis test with Dunn's multiple comparison test.

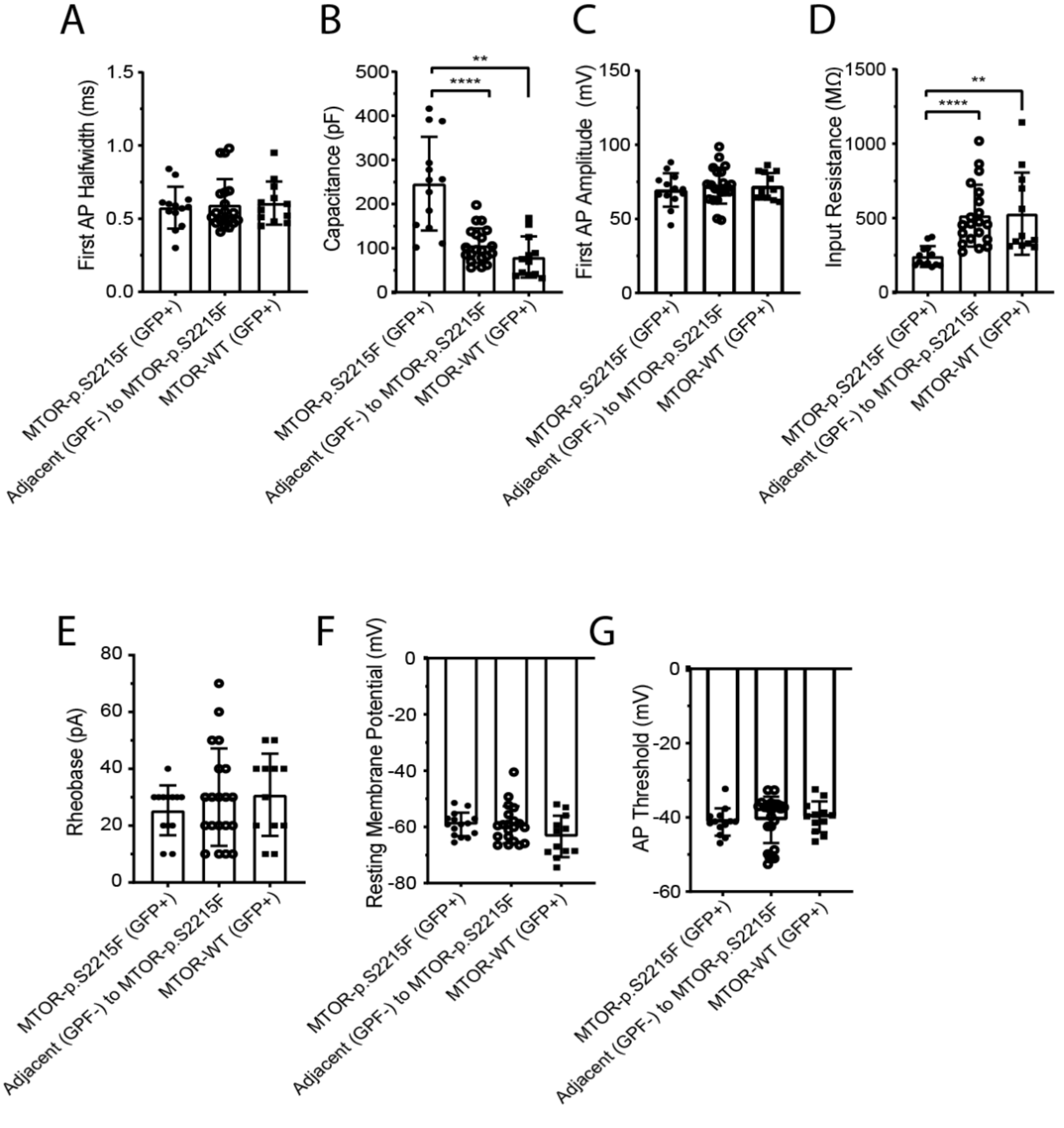

**Figure S5. Spontaneous EPSCs (sEPSCs) in electroporated and adjacent, non-electroporated neurons.**

Averaged and normalized to the peak sEPSCs were recorded in $Mg^{2+}$-free ACSF from MTOR-p.S2215F (grey, n = 10), MTOR-WT (red, n = 13) and pCAGIG-GFP (blue, n = 3) electroporated (GFP+) neurons, and from adjacent, non-electroporated (GFP-) neurons from MTOR-WT (purple, n = 7) and MTOR-p.S2215F (green, n = 2) electroporated pups aged P14-P16. Values are given as mean ± SEM. Note the larger area under curve of MTOR-p.S2215F sEPSCs compared to the other conditions.

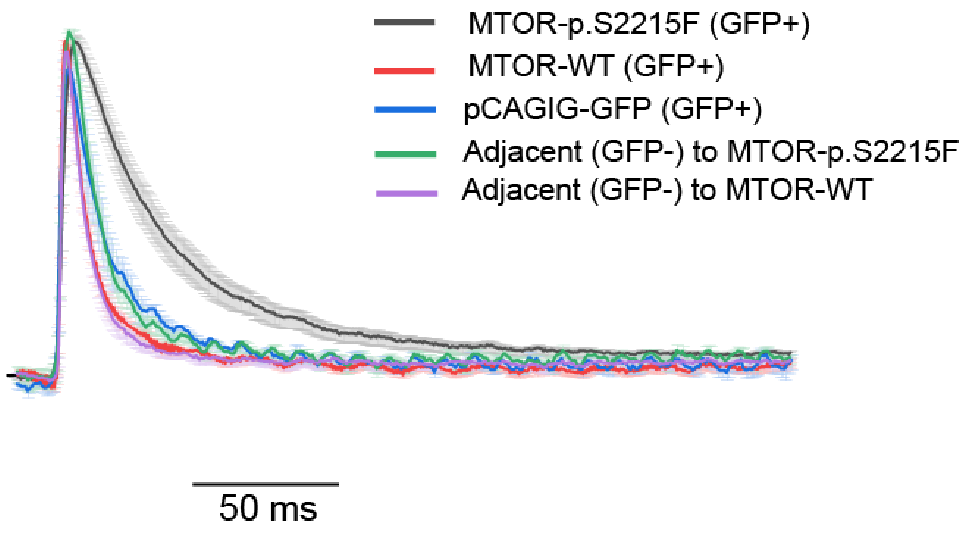

**Figure S6. No effect of DQP 1105 on spontaneous EPSCs (sEPSCs) in rapamycin-treated, MTOR-p.S2215F electroporated pups.**

(A) Representative traces of averaged sEPSCs recorded in MTOR-p.S2215F neurons from rapamycin-treated rats in (black curve): drug-free (No drug) condition, representing total component of sEPSC; (red curve): after adding GluN2C-mediated NMDAR blocker DQP 1105 (10 µM), representing the non-GluN2C-mediated component of sEPSC; and (blue curve): after subtraction of the non-GluN2C-mediated component from total sEPSC, hence representing GluN2C-mediated component of sEPSC (B) Charge transfers were calculated from MTOR-p.S2215F neurons in rapamycin treated animals as areas between peak and 200 ms after the peak of sEPSC with: No drug; GluN2C blocker DQP1105 (DQP; 10 µM); and total NMDAR blocker APV (50 µM). ns: not significant, Mixed effect analysis.

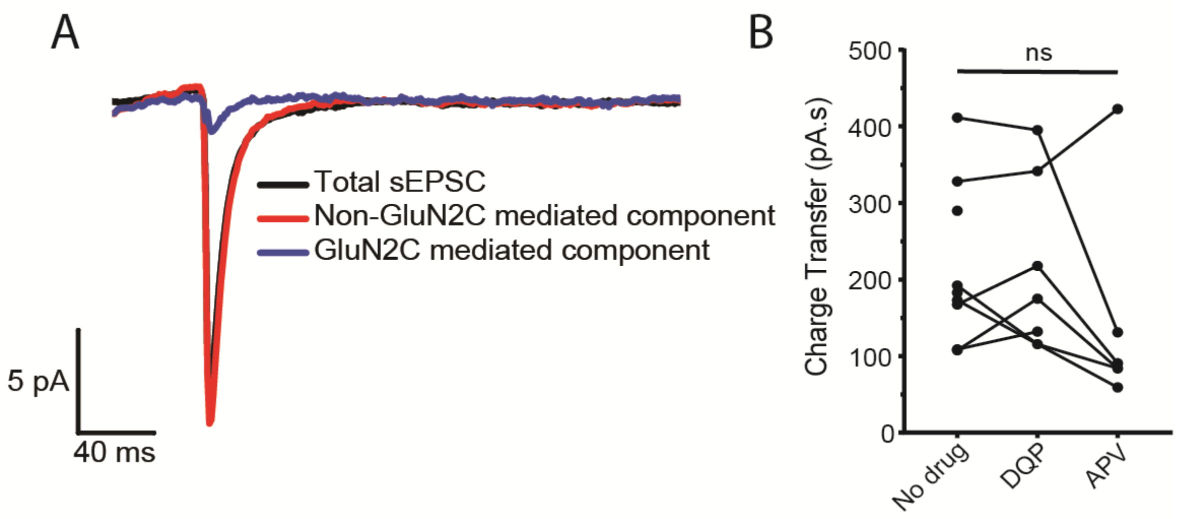

**Figure S7. GluN2C functional upregulation leads to increase synaptic temporal integration in MTOR-p.S2215F neurons.**

(A) Superimposed raw traces of spontaneous excitatory post-synaptic potentials (sEPSPs) recorded from MTOR-p.S2215F neurons in current clamp mode in whole cell configuration in $Mg^{2+}$-free ACSF without (left, n = 54 traces) and with (right, n = 69 traces) DQP 1105 (DQP, 10 µM). Note the higher number of spikes in the No drug condition (B) Average traces of sEPSPs recorded from MTOR-p.S2215F neurons in current clamp in $Mg^{2+}$-free ACSF without (left) and with (right) DQP. Note the larger area under curve in the No drug condition (see inset) (C) Quantification of the number of spikes for each sEPSP shown in (A) without (left) and with (right) DQP. Note the high number of 3 or 4 spikes in the No drug condition (time of recordings was 10 min for No drug condition and 14 min for DQP condition).

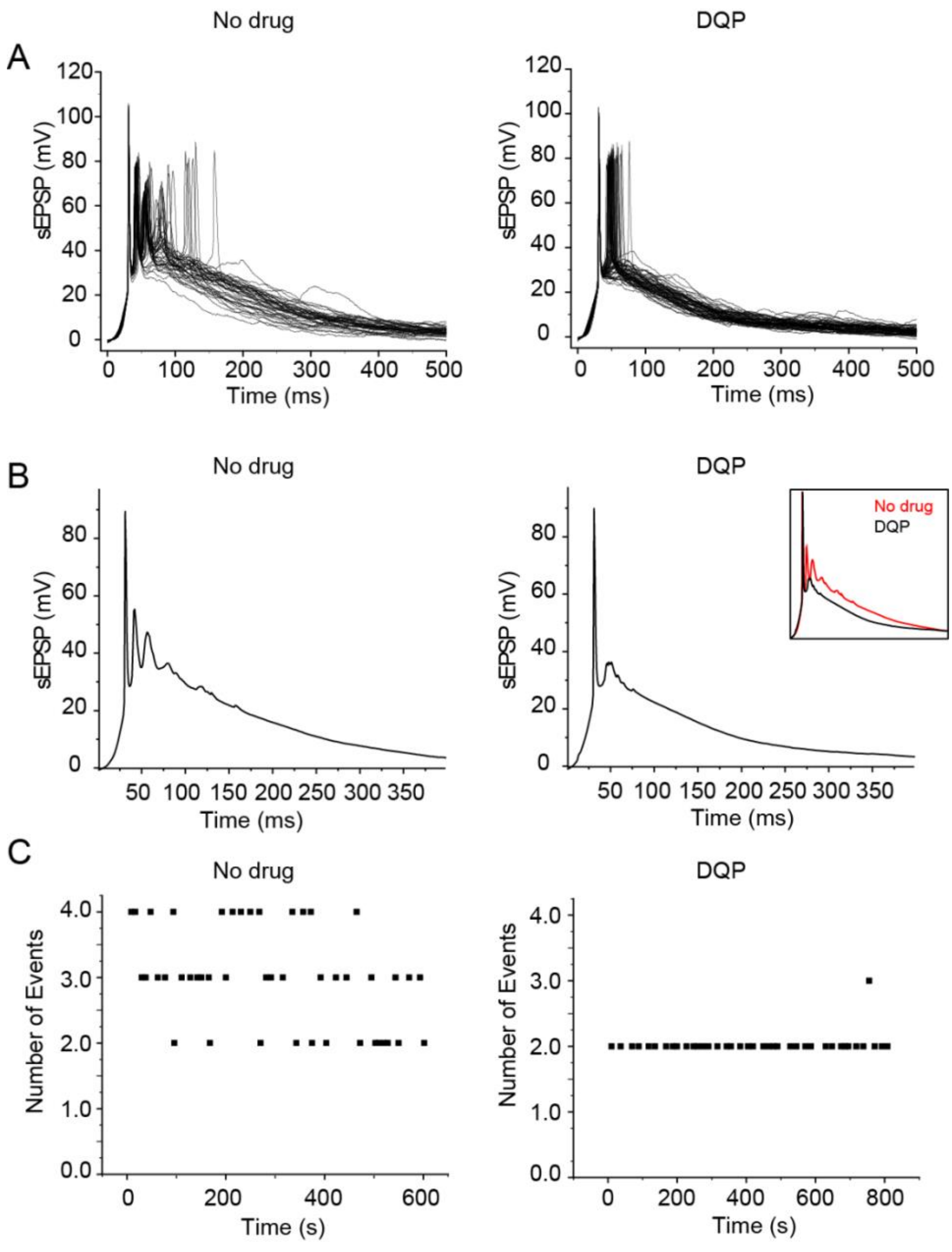

**Figure S8. Functional upregulation of GluN2C-containing NMDARs in mTOR p.S2215F electroporated neurons at P9.**

(A) Representative traces of sEPSCs from a neuron expressing MTOR-p.S2215F, recorded in $Mg^{2+}$-free ACSF at -75 mV without (No drug, left) or with 10 µM DQP 1105 (DQP, right) (B) Representative traces of averaged sEPSCs recorded in a MTOR-p.S2215F neuron in drug-free (No drug) condition (black curve), representing total component of sEPSC; after adding GluN2C-mediated NMDAR blocker DQP (10 µM) (red curve), representing the non-GluN2C-mediated component of sEPSC; and after subtraction of the non-GluN2C-mediated component from total sEPSC (blue curve), hence representing GluN2C-mediated component of sEPSC. sEPSCs of MTOR-p.S2215F electroporated neurons at P9 showed a strong DQP-sensitive component consistent with GluN2C-containing NMDAR-mediated currents.

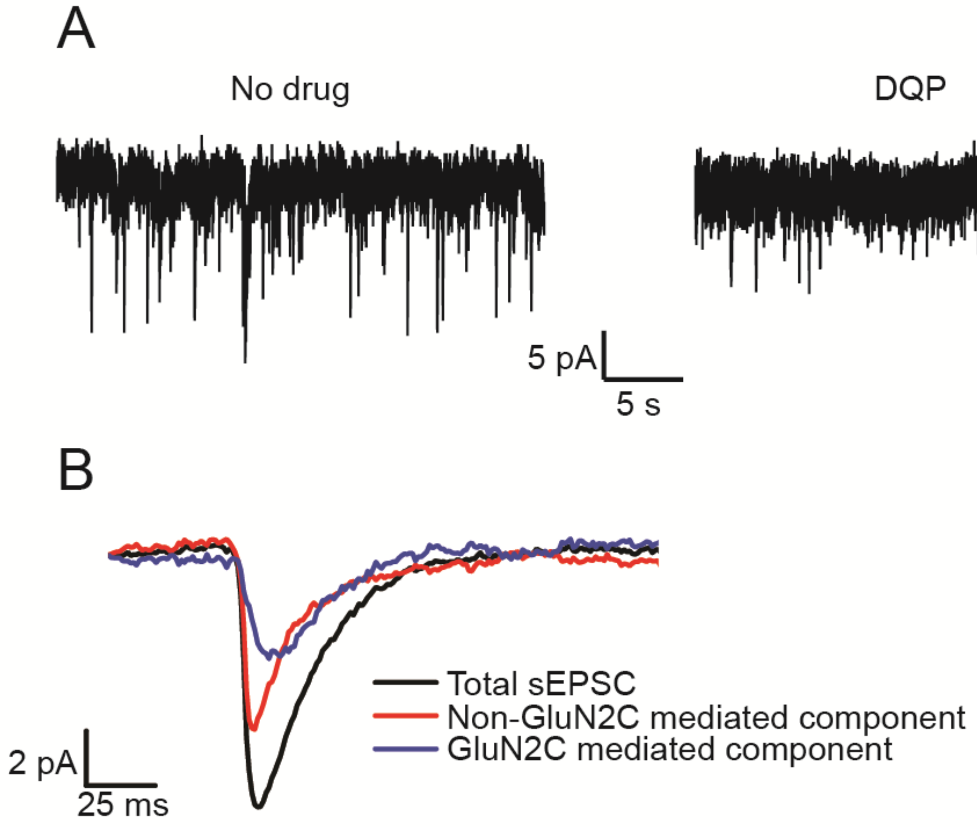
